# Genotype-environment-driven dysbiosis in the skin microbiome of ichthyosis

**DOI:** 10.1101/2024.06.17.599414

**Authors:** Wei Zhou, Nan G. Ring, Ryan Caldwell, Leonard M. Milstone, Julia Oh

## Abstract

Monogenic skin disorders such as ichthyosis introduce multiple sources of disturbance to the skin, including the direct biochemical consequences of the genotype, the phenotypic changes in skin physiology, and an altered skin microbiome. The association between changes in the skin microbiome and the disease’s genotypic and phenotypic effects are of both ecological and clinical interest but are historically obscured by 1) the limited resolution of metagenomic profiles, and 2) additional sources of variation such as age and topical/oral treatments. Here we characterize the skin microbiome from seven ichthyosis genotypes, at species, strain, and metabolic pathway levels. Critically, we assess the association between these microbiome features and the ichthyosis genotype and phenotype while adjusting for contextual host covariables. We show that the ichthyosis genotype, especially that caused by mutations in TGM1, and the ichthyosis phenotype, particularly transepidermal water loss (TEWL), and personal covariables, such as topical emollients and oral retinoids, collectively, and sometimes antagonistically, influence the species community, strain population, and metabolic potential of the skin microbiome.

**One Sentence Summary:** Species, strain, and the composition of metabolic pathways of the cutaneous microbiome in ichthyosis are influenced by the mutation genotype of ichthyosis, phenotypic features of ichthyosis such as transepidermal water loss, scaling, and skin hydration, and host age and treatment.

## INTRODUCTION

The remarkable diversity of the bacterial, fungal, and viral species comprising the human skin microbiome has critical roles in skin barrier maintenance (*1*), immune modulation (*2, 3*), and protection against pathogens (*4*). This diversity is strongly shaped by the type of skin site, with oil-rich skin sites (e.g., face and torso) predominated by *Cutibacterium* bacteria and *Malassezia* fungi, compared with a relatively higher diversity of Actinobacteria, Firmicutes, and Proteobacteria in dry sites (e.g., hands and forearm) or moist sites (e.g., armpit) (*5, 6*). Such environment-dependent distribution of microbial taxa is also hypothesized to vary due to differences in host genetics (*7, 8*). On top of that, the complex associations between host genetics and the skin microbiome have been difficult to unravel due to numerous covariables, including the physiological conditions of the skin, the age of the subject, and the use of therapeutic or cosmetic products. The overwhelming majority of skin microbiome studies to date have not adjusted for such covariables, complicating the interpretation of differences between cohorts and the generation of testable hypotheses.

Here, we assessed the host genetics-microbiome association by performing a high-resolution compositional and functional reconstruction of the skin microbiome with a rigorous adjustment for covariables. To highlight the influence of host genetics on the microbiome, we examined the skin microbiome of patients with monogenic mutations causing ichthyosis. The ichthyoses are a family of disorders leading to widespread dry, scaly, peeling, or thickened skin (*9*). Notably, most cases of ichthyosis are Mendelian, caused by mutations in a wide variety of genes, resulting in impaired keratinocyte differentiation and skin barrier dysfunction (*9, 10*). In normal skin, the most important component of the skin barrier is the stratum corneum, which consists of dead cells filled with tightly packed keratin filaments surrounded by an envelope of cross-linked protein and extracellular lipids produced by keratinocytes and secreted by the lamellar bodies. Correspondingly, mutations in genes coding for proteins involved in cornified cell envelope formation (e.g., *TGM1*), lipid metabolism or secretion (e.g., *ABCA12*, *ALOX12B*, and *NIPAL4*), or keratin filaments (e.g., *KRT1*, *KRT2*, and *KRT10*) cause ichthyosis and disrupt normal barrier function. Ichthyosis can also be caused by mutations in *SPINK5*, disrupting the barrier primarily by allowing the stratum corneum to slough prematurely.

The phenotypic consequences of the genetic mutations of ichthyosis include a dysfunctional stratum corneum characterized by increased transepidermal water loss (TEWL), decreased skin hydration, and scaling (*9–11*). Taken together with the monogenic disorder-specific biochemical changes in the skin, these phenotypic factors inevitably affect the skin microbiota, which is likely responsive to pathological changes in the skin condition (*12, 13*). It is also conceivable that ichthyosis-associated dysbiosis of the skin microbiome could reciprocally exacerbate the disease state, and some types of ichthyosis have increased susceptibility to skin infection. From a mechanistic perspective, identifying which aspects of ichthyosis the skin microbiome responds to (e.g., the direct biochemical outcome of the genotype or measurable phenotypic changes such as TEWL, altered level of hydration, or the severity of scales) is necessary for generating testable hypotheses. Moreover, the correlations between ichthyosis genotype or phenotype and skin microbes are inevitably convoluted by other characteristics of the host that alter the skin microbiome. For example, ichthyosis patients commonly receive topical and oral treatments, which can alter the skin microbiome (*14–16*). In addition, we have found that the skin microbiome is famously personalized and age-dependent (*17, 18*).

In view of these considerations, our overarching goal was to determine whether the genotypic effect of ichthyosis – the direct biochemical changes specific to the monogenic disorder – and measurable phenotypic features of the stratum corneum that these monogenic disorders have in common—increased TEWL, decreased skin hydration, increased scaling—impact the composition of the local skin microbiome. Characterizing the skin microbiome in ichthyosis would provide the necessary foundation for understanding the basic mechanistic interactions between skin substrates provided by the host and the skin microbiome. In turn, this information is necessary for understanding the role of the microbiome in skin health both in healthy and ichthyotic skin, potentially expanding the repertoire for clinical solutions for skin infection risk in these patients via modulation of the cutaneous microbiome.

Despite recent efforts (*13, 19*), a detailed and contextualized profile of the ichthyosis skin microbiome is still lacking. This is partly due to the variety of ichthyosis-causing mutations and the diversity in the skin microbiome itself: variations in the skin microbiome are often manifested at the species, strain, or gene level (*6, 20–23*). Studies to date have used low-resolution approaches such as amplicon sequencing to profile the ichthyosis microbiome (*12, 19*), or have only examined species composition at a limited number of skin environments (*13*), while the drastically different microbiome compositions at dry, oily, moist, and foot environments could lead to niche-specific ichthyosis-microbe interactions. Finally, none have explicitly accounted for genotypic, phenotypic, and host covariables.

Here, we present the first high-resolution and contextualized profiling of the ichthyosis skin microbiome, representing seven ichthyosis-causing genotypes, examining ten skin sites, and integrating phenotypic, host age, and clinical treatment information. Strikingly, we not only observed a significant variation in microbiome diversity between individual genotypes but were also able to identify genotype-associated microbes. Using TGM1 as the focal genotype, we were able to attribute variations in the microbial community to genotypic effects, phenotypic effects, and other host features. We found that, at the species level, the ichthyosis microbiome composition was primarily dependent on both the genotypic effect and the phenotypic variable TEWL, with the two variables often acting antagonistically. We also found that Aquaphor treatment, specifically on TGM1 skin, contributed to a robust enrichment of *Corynebacterium (Co.) resistens* in the TGM1 patients. At the strain level, although the TGM1 genotype appeared a major determinant of the *Co. resistens* strain population, TEWL had the most significant effect on the structures of the strain populations of other abundant species, highlighting how stratum corneum integrity affects the microbiome at both community and population level. Additionally, a dysfunctional stratum corneum also appeared to select for microbes scavenging multiple biological compounds from the environment, as opposed to microbes synthesizing them *de novo*, potentially due to the increased availability of these compounds in the damaged skin. Taken together, this study showed that the ichthyosis genotype, phenotype, and personal covariables collectively influence the species community, strain population, and metabolic potential of the skin microbiome.

## RESULTS

### Study Design

Our goal in this study was to identify changes in the microbiome in ichthyosis and demonstrate whether they were driven by the genotype, phenotype, or other covariables – treatment and host age. The rarity of many of the individual ichthyosis genotypes, other than TGM1, precluded our recruiting enough individuals with single genotypes to correlate microbiome features simultaneously with differences in skin hydration, TEWL, and the quality and severity of scale between single genotypes of ichthyosis (Table S1 and Fig. S1) and the well-described biogeographic diversity in the skin microbiome. Therefore, we first conducted a single variable screening to show that, when analyzed individually, phenotypic variables and microbiome features do manifestly differ across ichthyosis genotypes. (Hereafter, we will use the word “manifest” to describe the observed differences between genotypes *without* adjusting for phenotypic effect or host covariables). Second, we based the bulk of our analyses focusing on the TGM1 genotype while adjusting for phenotypic effect and host covariables. We do this by grouping the other ichthyosis genotypes into “non-TGM1 ichthyosis” (NTI) to contrast against TGM1. At the same time, we grouped the ten sampled skin sites into four skin types: dry (upper arm, shin), moist (intergluteal cleft, popliteal fossa), oily (scalp vertex, forehead, central chest, upper back, postauricular), and sole based on known similarities in their microbiota content (*6*), then hierarchically modeled the variation between the skin sites within each skin type as random effects. Additionally, we present a parallel analysis while grouping the non-TGM1 ichthyosis genotypes by their direct effect on critical components of the stratum corneum: keratins, lipids, and cornified envelopes (Supplementary Note). The keratinopathic ichthyoses (KPI) are represented KRT2 and KRT10; lipid deficient ichthyoses (LDI) are represented by ABCA12, ALOX12B, and NIPAL4. SPINK5 samples were not included in this analysis because of their singular mechanism and limited sample size.

### Skin microbiome and phenotypic features manifestly differ across ichthyosis genotypes

We first screened the ichthyosis samples for genotype-specific microbiome patterns, hypothesizing that the skin microbiome composition was not only manifestly different between ichthyosis patients and healthy controls but also manifestly different between patients of distinct ichthyosis genotypes. Indeed, the healthy controls exhibited a consistent dominance of *C. acnes* at the oily sites, while the ichthyosis skin had more abundant *Corynebacterium* and *Staphylococcus* bacteria (Fig. 1A). Three ichthyosis patients (ICHT-2, ICHT-4, and ICHT-46) had a substantial fraction of metagenomic space that failed to be classified by MetaPhlAn4 (*24*) (Fig. 1A), indicating overgrowth of previously uncharacterized microbes in a small proportion of the patients, or the presence of strain-level diversity in classified microbial species that were captured by the sequencing reads but not in public databases. For consistency of the results, we focused our initial downstream analyses on the classified microbes. Consistent with our hypothesis, we found a significant difference between the alpha diversity of the healthy skin microbiome and the alpha diversity of the ichthyosis microbiome at the oily sites and the moist sites (Fig. 1B and Table S2). Interestingly, compared to the ichthyosis microbiome, the healthy skin microbiome at the oily sites was significantly less diverse at the oily sites, but significantly more diverse at the moist sites (Fig. 1B and Table S2). This was likely because the lipophilic *Cutibacterium (C.) acnes* dominated the oily sites of the healthy skin (*6*) (Fig. 1A), which numerically decreased Shannon’s index. Such dominance was often lacking at the ichthyotic skin and the moist sites of the healthy (Fig. 1A) and may be due to decreased sebum secretion in those environments. We also found that the alpha diversity differed between ichthyosis genotypes at dry, oily, and sole sites but not at the moist sites (Table S3). Of the seven genotypes, the TGM1 microbiome was among the most diverse numerically at the dry, oily, and sole sites (Fig. 1B), although most of the pairwise comparisons of Shannon’s index between genotypes were not statistically significant (Table S4). In addition to the within-sample alpha diversity, the structures of the microbiome between healthy controls and ichthyosis patients and between ichthyosis patients of different genotypes also differed significantly in all skin environments (distance-based redundancy analysis based on Bray-Curtis distance p < 2.2×10-6), again showing that different ichthyosis genotypes can have distinctive influences on the skin microbiome.

**Fig. 1.**
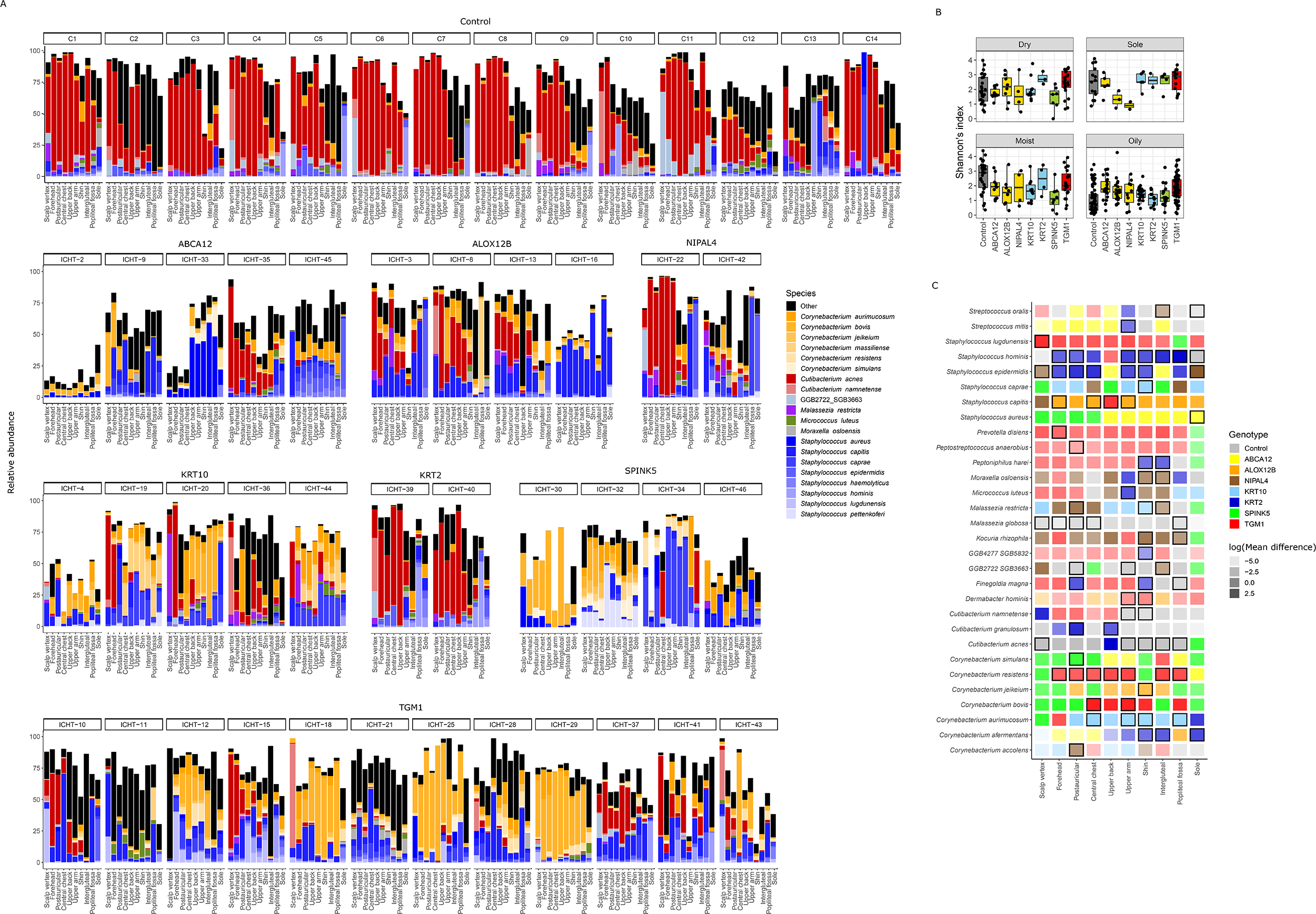
Species diversity of the ichthyosis microbiome. (**A**) Distribution of the top twenty most abundant microbial species. (**B**) Distribution of species-level alpha diversity as measured by Shannon’s index (N=48). P or q values of statistical comparisons were summarized in Table S2-S4. (**C**) Manifest enrichment of microbial species in ichthyosis genotypes. Transparency of the tile colors represents the mean difference in species relative abundance in one ichthyosis genotype (or the healthy control) versus all other samples. Tiles with black outlines represent species where the lower bound of the 95% confidence interval of the mean difference was greater than zero.

**Fig. 2.**
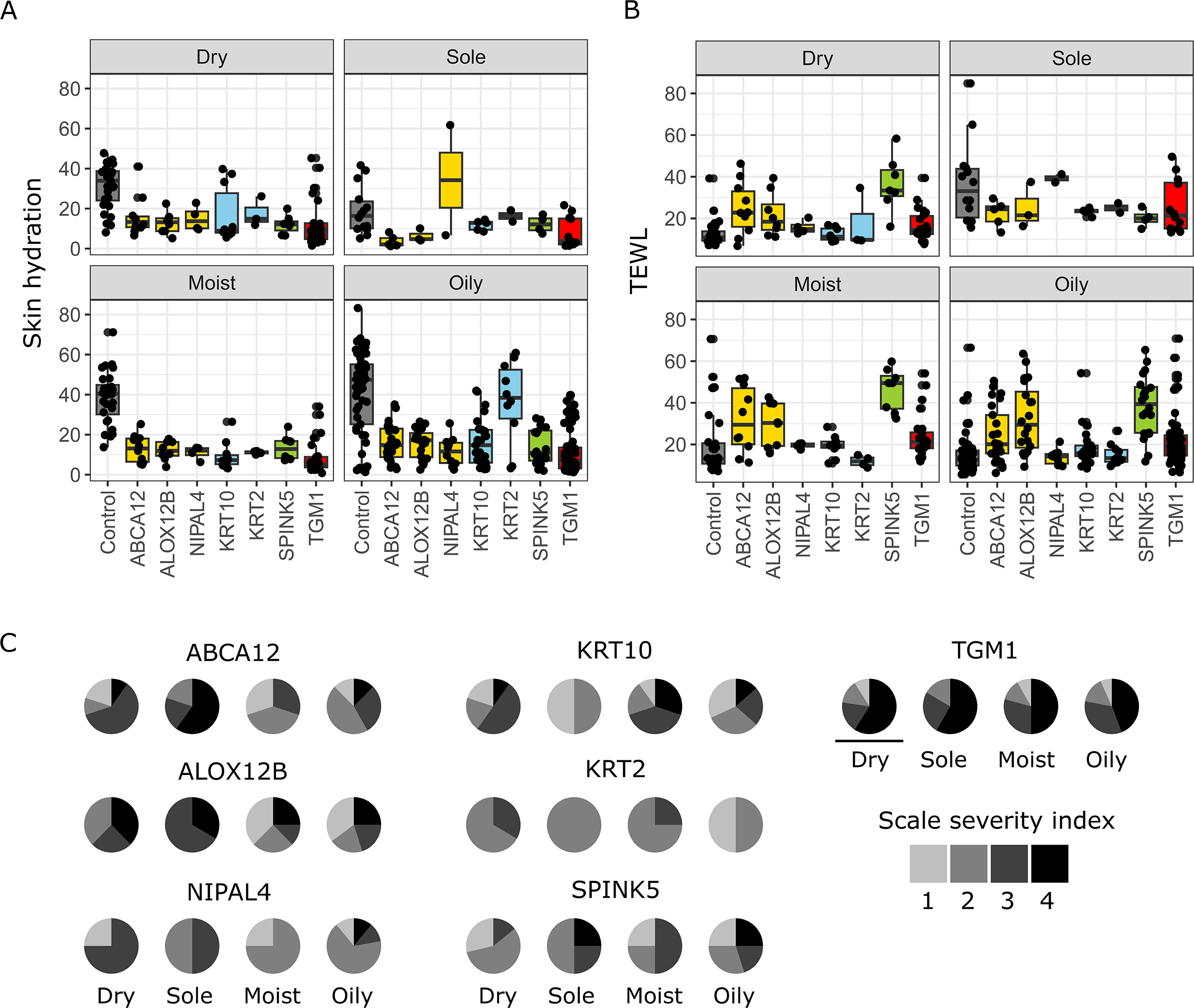
Distribution of ichthyosis phenotypes across ichthyosis genotypes. (**A**) Distribution of skin hydration (corneometer measurements) across ichthyosis genotypes. (**B**) Distribution of TEWL (tewameter measurements) across ichthyosis genotypes. (**C**) Distribution of scale severity index across ichthyosis genotypes. P or q values of statistical comparisons were summarized in Table S5-S9.

Next, we sought to identify those species that were distinctively and manifestly associated with each ichthyosis genotype. We used a nonparametric bootstrap method to identify microbial species that were enriched in one genotype, relative to all other genotypes, with at least a 95% confidence. As expected, *C. acnes* and *Malassezia (M.) globosa* were both enriched in the healthy controls at multiple body sites, representing the hallmarks of a healthy skin microbiome (Fig. 1C). Most surprisingly, *Co. resistens* was enriched in TGM1 at seven out of the ten skin sites with at least a 95% confidence (Fig. 1C), indicating adaptation to a genotype-specific condition. Other skin bacteria such as *Staphylococcus (S.) lugdunensis, Prevotella (P.) disiens,* and *Corynebacterium bovis*, also tended to be associated with TGM1 at most of the skin sites, although with lower confidence (Fig. 1C). We also observed that *S. capitis* tended to be enriched in ALOX12B at eight out of the ten skin sites, among which the enrichment at the forehead and upper arm met the 95% confidence threshold (Fig. 1C). While *S. epidermidis* and *S. hominis* did show enrichment in KRT2 patients at multiple sites, the result could be due to an inadequate representation of the KRT2 diversity (patient N=2). Other microbe-genotype associations appeared more nuanced and skin site-specific. For example, *M. restricta* was enriched in the NIPAL4 microbiome at the postauricular and intergluteal sites but was enriched in the KRT10 microbiome at the shin (Fig. 1C). We also observed similar results when ichthyosis genotypes were grouped based on their functional mechanisms (Supplementary Note Fig. 1A).

These manifest microbiome differences observed between ichthyosis genotypes were inevitably influenced by the disease phenotype and host covariables. To deconvolute the multifaceted relationships, we sought to characterize the phenotypic manifestation of each ichthyosis genotype on the skin. We found significant differences in skin hydration and TEWL measurements between healthy controls and all ichthyosis patients (Fig. 1A, 1B, and Table S5). Among ichthyosis patients of different genotypes, the skin hydration at the oily sites and the sole, as well as TEWL at all environments except for the sole were also significantly different (Table S6), strongly indicating that phenotypic conditions of the skin were genotype-dependent. Numerically, SPINK5 tended to show the highest TEWL measurement in all skin environments except for the sole (Fig. 1B), consistent with a previous study (*25*). A post hoc pairwise comparison revealed that a substantial proportion of the trend was statistically significant (Table S7). TEWL had been shown to be positively and manifestly correlated with the relative abundance of *Corynebacterium* and *Staphylococcus* in healthy individuals (*26*); while we found a positive and manifest correlation between TEWL and the relative abundance of the *Staphylococcus* genus in the healthy control (Pearson’s *r*=0.25, p=0.0029), no significant correlation was observed between TEWL and the relative abundance of the *Corynebacterium* genus (Pearson’s *r*=-0.0094, p=0.91). In terms of skin hydration, TGM1 skin trended towards the most dehydrated among all genotypes (Fig. 1A), although the pairwise differences were mostly statistically insignificant (Table S8). The distribution of the scale severity index also significantly differed across ichthyosis genotypes at all skin environments except for the sole (Fig. 1C and Table S9). In terms of scale severity, the genotype that had the most extreme phenotype was again TGM1, which consistently exhibited the largest proportion of samples with level 4 scale severity across skin environments (Fig. 1C). In addition to the skin phenotypes, we also looked at the distribution of host covariables across genotypes. We found that, although the age of the patients did not significantly differ across genotypes (ANOVA p=0.98), both topical treatments did (Chi-squared test p=0.04 for Vaseline and p=0.0038 for Aquaphor) and only TGM1 patients received retinoid treatment.

### Microbial diversity in TGM1 skin is modulated by stratum corneum integrity and topical treatment

Thus far we observed manifest enrichment of microbial species in distinct ichthyosis genotypes, but it was unclear if these enrichments were due to the direct biochemical changes induced by the ichthyosis genotype, measurable phenotypic changes that alter the skin environment the microbes lived in, or other host characteristics. We sought to differentiate these factors, hypothesizing that microbial species variations could be explained by genotypic and phenotypic changes of ichthyosis as well as covarying factors such as host age and treatments. To approach this question, we conducted two analyses, comparing TGM1 samples to healthy controls (TGM1+healthy control models) and non-TGM1 ichthyosis (NTI) samples (TGM1+NTI models), respectively. To account for the substantial biogeographic variation in the skin microbiome, we analyzed the dry, moist, and oily skin sites separately. Sole samples were not included in these analyses because no TGM1 patients had a scale severity index equal to 1 at their sole (Table S1).

We first investigated what variables – genotypic, phenotypic, or other host covariables – influenced the microbiome alpha diversity. By modeling the genotypic, phenotypic, and confounding variables using a mixed effect model, we found that the observed variation in alpha diversity was not only attributable to the TGM1 genotype but also dependent on TEWL (Fig. 3A, Table S10, and Table S11). Interestingly, even without adjusting for other variables, increased TEWL demonstrated an overall negative association with Shannon’s index, except for at the moist sites in TGM1 (Fig. 3B). The other phenotypic variables and host covariables were not significantly associated with alpha diversity. These findings suggested potentially competing factors driving alpha diversity in skin: the genotypic effect of TGM1 increased local alpha diversity, while its disruption of the stratum corneum dampened alpha diversity. More broadly speaking, our findings demonstrated that both genotypic and phenotypic differences influence the skin microbiome.

**Fig. 3.**
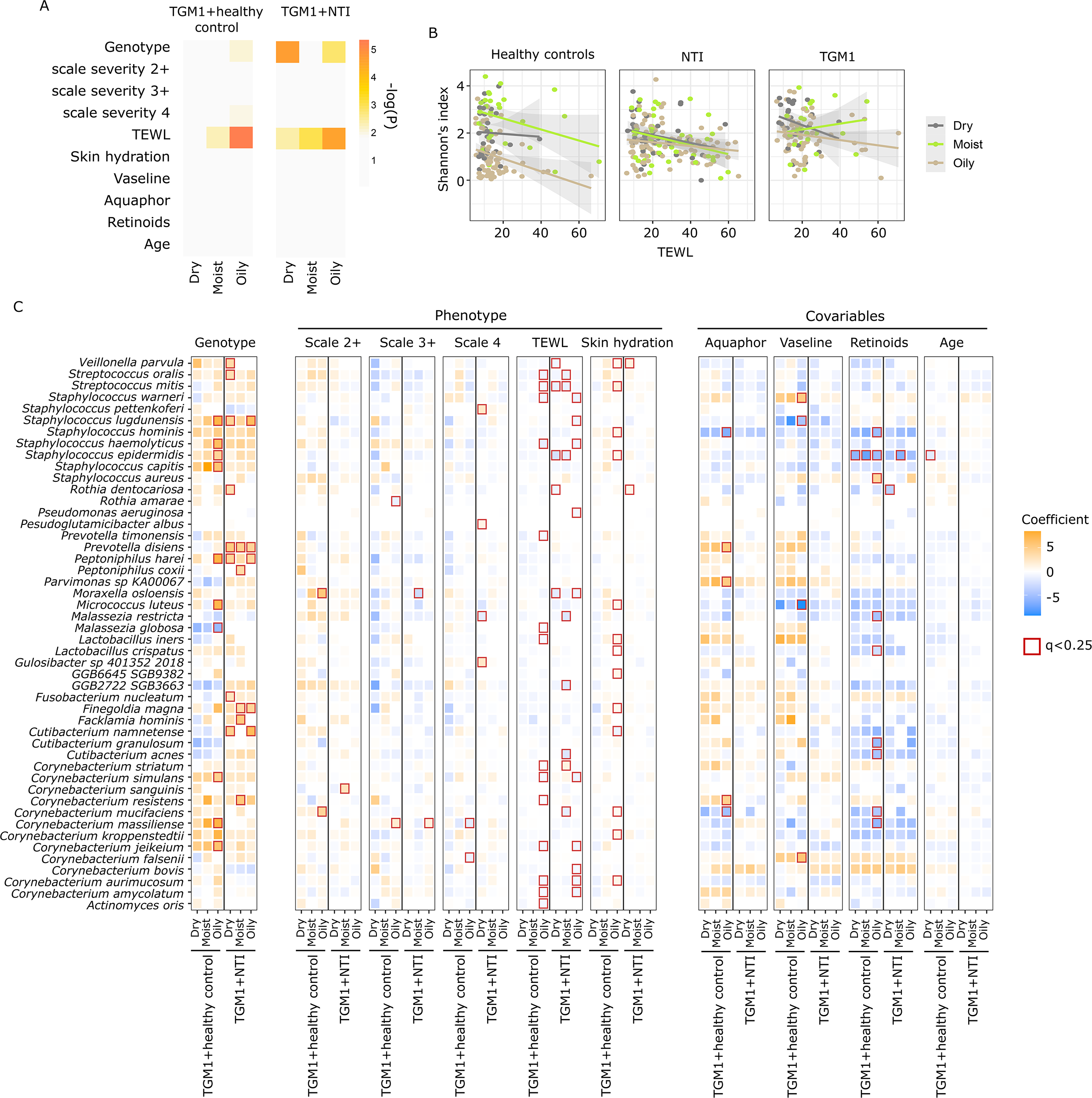
Associations between microbial species and host variables. (**A**) Significance of associations between species Shannon’s index and host features. P-values used are summarized in Table S10 and S11. (**B**) Relationship between TEWL and Shannon’s index without adjusting for other host variables. The lines represent linear regressions between TEWL and Shannon’s index, and the shaded areas represent 95% confidence interval. (**C**) Associations between species relative abundances and host variables estimated using MaAsLin2. Significant associations (q < 0.25) had red outlines. Only species with at least one significant association were shown. Values used for this figure were summarized in Table S12.

Given this multifactorial influence on the ichthyosis microbiome, we next asked which genotypic, phenotypic, and host covariables are associated with each of the microbiome species. We tested multivariable associations between the microbial species abundance and the heterogeneous genotypic, phenotypic, and covarying variables using MaAsLin2 (*27*). Similar to what was observed for Shannon’s index, we found that genotype and TEWL influenced the skin microbiome most significantly in both the TGM1+healthy controls models and TGM1+NTI models (Fig. 3C and Table S12). Skin hydration also significantly correlated with the relative abundance of multiple skin species, but these effects were mostly observed at the oily sites in the TGM1+healthy control model. Interestingly, many of these hydration-associated species, including *S. hominis, S. epidermidis*, *Micrococcus* (*M.) luteus, Cutibacterium namnetense, Finegoldia magna, Corynebacterium mucifaciens,* and *Corynebacterium kroppenstedii*, were not significantly affected by TEWL at the oily sites. On top of the genotypic and phenotypic influence, medical treatment also altered microbial relative abundances, while the effect of host age was insignificant on most species (Fig. 3C and Table S12). This interaction of genotypic, phenotypic, and treatment effects was observed for several species that were shown to be manifestly enriched in TGM1. For example, a robust association was observed between *S. lugdunensis* and TGM1, which was statistically significant at the oily sites and moderately positive at the dry and moist sites compared to both the healthy controls and NTI (Fig. 3C and Table S12). Contrary to the genotypic effect, Vaseline treatment and increased TEWL at the oily sites tended to decrease *S. lugdunensis* abundance. Similarly, the robust manifest enrichment of *Co. resistens* observed in TGM1 across almost all body sites (Fig. 1C) could be explained by the genotypic effect and the application of medical treatments (mainly Aquaphor) collectively (Fig. 3C and Table S12). Interestingly, the association between *Co. resistens* relative abundance and Aquaphor treatment was only positive in the TGM1+healthy control models and only significant in oily skin (Fig. 3C and Table S12). Because these treatments were not applied to healthy controls, the TGM1+healthy control models can only assess the effect of the Aquaphor treatment by comparing TGM1 samples with or without the treatments. Therefore, we deduce that the positive association between *Co. resistens* abundance and Aquaphor treatments was only present in TGM1 patients, suggesting a genotype-specific treatment effect. On the contrary, the enrichment of *Co. resistens* in TGM1 compared to NTI appeared to be mainly driven by the genotype effect but not Aquaphor treatments (Fig. 3C and Table S12). Similarly, *Prevotella (P.) disiens*, which also showed a tendency of enrichment in TGM1, was positively associated with the TGM1 genotype in TGM1+NTI models, while positively associated with the Aquaphor treatment in the TGM1+healthy controls models (Fig. 3C and Table S12). Consistent with a study on atopic dermatitis (*16*), we observed a tendency of increase in the relative abundance of *C. acnes* and *M. globosa* with Vaseline and Aquaphor treatment, but these associations were not significant (Fig. 3C and Table S12). As retinoid treatment in this study was only applied to TGM1 patients, its effect in TGM1+healthy control and TGM1+NTI models were highly consistent, with the most significant change observed in *S. epidermidis*, whose relative abundance decreased with retinoid treatment. We also observed that retinoids decreased the relative abundance of *Cutibacterium* species especially at the oily sites – a pattern previously found in the acne skin (*15*). Taken together, by statistically attributing observed variation to the underlying factors, our multivariable analysis revealed how the effect of the TGM1 genotype, TEWL, and treatment collectively resulted in the observed enrichment of *S. lugdunensis, S. epidermids, Co. resistens*, and *P. disiens* in the TGM1 patients.

### Strain-level diversity is associated with TEWL and lacks phylogenetic signal

One possibility for the observed complexity of genotypic, phenotypic, and covarying effects on species relative abundance is that there exists strain-specificity in these interactions. For example, the observed enrichment of *Co. resistens* in TGM1 could be driven by a few phylogenetically close *Co. resistens* strains (a pattern observed in *S. aureus* strain enrichment in atopic dermatitis, which is primarily clonal), or it could be the result of a relatively uniform enrichment of all *Co. resistens* strains (like heterogeneous *S. epidermidis* strain diversity observed in atopic dermatitis) (*28*). To explore these possibilities, we leveraged the resolution provided by shotgun metagenomic data to ask if there were strain-level variations associated with genotype, phenotype, and covariables. We performed *de novo* assembly and metagenomic binning to generate 907 non-redundant metagenome-assembled genomes (MAGs) from the samples used in this study (Fig. S2), which we then used to delineate the strain abundances of these samples. 365 of our 907 MAGs had >90% completeness and <5% contamination; the remainder had >50% completeness and 10% contamination (Table S13). The advantage of using MAGs over public genome databases is that MAGs delineate strain-level differences that are specific to this dataset, and thus it is more likely to uncover strain features and associations specific to this dataset using MAGs rather than a public genome database. For an adequate and accurate representation of the strain diversity, we focused on four species (*C. acnes*, *Co. resistens*, *Co. amycolatum*, and *S. capitis*) that 1) were each represented by at least 15 MAGs, and 2) had a total strain abundance close to the estimated species abundance (Pearson’s *r*>0.95, Table S14), supporting that the MAGs accounted for most of that species’ strain variation. As with species-level analyses, we modeled multivariate associations between MAG abundances and genotype, phenotype, and covariates.

Compared to the species-level analyses, we found fewer significant associations between genotype and MAGs (Fig. 4A and Table S15-S18), suggesting a less prominent genotypic effect in structuring the strain population compared with structuring the species community. An exception to this is the enrichment of eight *Co. resistens* MAGs at the TGM1 oily sites compared to the healthy controls, with two of the eight also enriched at the TGM1 dry sites compared to the healthy controls (Fig. 4A and Table S16). These MAGs also tend to show higher relative abundances in TGM1 compared to NTI samples, but the difference was not statistically significant (Fig. 4A and Table S16). Similar to the genotype effect, the influence of treatments and age on the strain population was also nuanced and rarely significant. On the contrary, the phenotypic variable TEWL negatively correlated with a large array of MAGs in all four species analyzed (Fig. 4A and Table S15-S18). These correlations were mostly observed at the moist or oily sites in both the TGM1+healthy control models and the TGM1+NTI models, suggesting that, a dysfunctional stratum corneum characterized by increased TEWL tends to suppress strain diversity at moist and oily skin sites, at least for the four species analyzed. Given that increased TEWL also suppressed diversity at the species resolution (Fig. 3B and 3C), we conclude that stratum corneum dysfunction can cause dysbiosis across taxonomic levels.

**Fig. 4.**
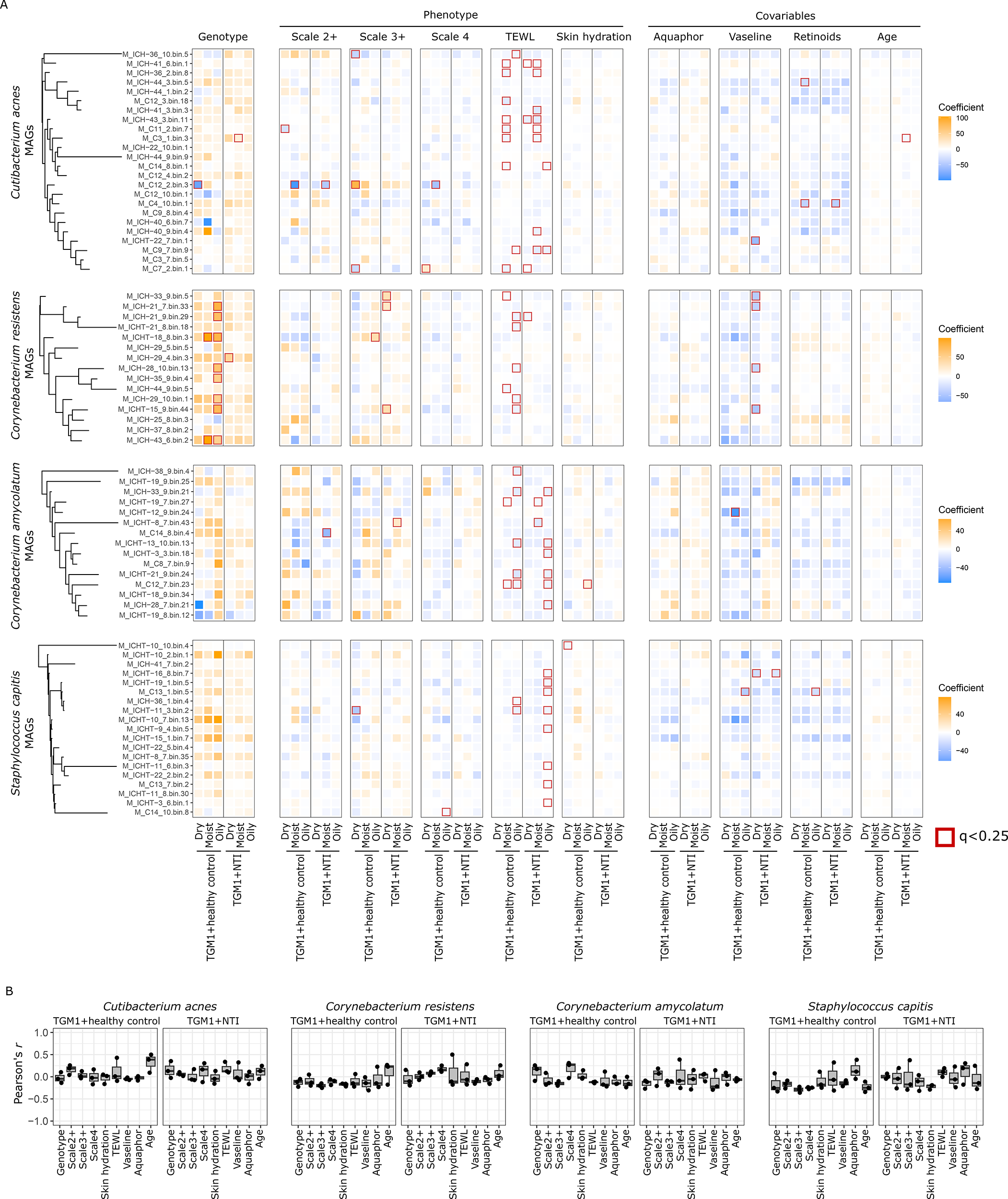
Associations between microbial strains and host features. (**A**) Associations between the relative abundances of MAGs from four abundant species and host variables estimated using MaAsLin2. The relative abundances of the MAGs were rescaled within each species to sum up to 1. Phylogenetic relationships of the MAGs from each of the four species were illustrated using a subtree of Figure S2. Significant associations (q < 0.25) had red outlines. Values used for this figure were summarized in Table S15-S18. (**B**) Pearson’s correlation coefficient between cophenetic distance of MAGs and the MAGs’ differential response to host variables. Each data point represents a skin environment (dry, oily, or moist).

Because most of the associations between MAG abundances and the host variables (other than TEWL) were MAG- and skin environment-dependent, we asked what aspects of the MAGs determined such nuanced associations. One hypothesis was that phylogenetically closer MAGs associated with genotypic, phenotypic, and other covariables in more similar ways. We tested this hypothesis by asking if the cophenetic distance between a given pair of MAGs – a proxy for their phylogenetic difference – was positively correlated with the difference in their MaAsLin2 coefficient for a given variable – a proxy for the difference in the MAGs’ response to changes in that variable. Surprisingly, we found that the phylogenetic similarity of MAGs was generally not positively correlated with the MAGs’ associations with genotypic, phenotypic, and host covariables (Fig. 4B). One exception to this general lack of phylogenetic signal was that phylogenetically close *C. acnes* strains changed with age in more similar ways than phylogenetically distant strains in the TGM1+healthy control models (Fig. 4B). Interestingly, we found that the correlation coefficients between phylogenetic differences and the differences in associations with host variables tended to be slightly negative in the TGM1+healthy control models, especially for *S. capitis* MAGs (Fig. 4B). This suggested that phylogenetically close *S. capitis* strains could experience more antagonistic interactions due to, for example, competition for similar types of resources, than phylogenetically distant strains. Although our present dataset does not have enough resolution to address these hypotheses, our findings do suggest that in the ichthyosis microbiome, 1) a strain’s response to host variables could be influenced by features other than its phylogenetic position, and 2) a more comprehensive characterization of the strain-level diversity using methods such as large-scale isolate sequencing is required to identify such features.

### Abundance of various biosynthetic pathways is altered in ichthyotic skin

One of the microbial features that does not segregate along phylogenetic or taxonomic boundaries is the microbial metabolic pathways; the observed association between microbial species and strains with ichthyosis could be a result of natural selection on the metabolic pathways carried by these microbial species and strains. Therefore, we sought to identify microbial metabolic pathways associated with host genotype, phenotype, and covariables. To focus on pathways that were robustly associated with host variables, instead of pathways that were merely hitchhiking other genomic loci under selection, we reported pathway associations not only significant on the community level but also concordant at the species level (detailed in the Materials and Methods section). With this strict filter we were only able to identify significant associations at the moist and oily sites. We found that the TGM1 genotype enriched for biosynthetic pathways of branched-chain amino acids (Metacyc ID: PWY-5103, ILEUSYN-PWY, BRANCHED-CHAIN-AA-SYN-PWY) at both the oily and the moist sites when compared to healthy controls, but not when compared to NTI samples (Fig. 5 and Table S19), suggesting that the branched-chain amino acids biosynthetic pathways were enriched in not just TGM1 skin, but ichthyotic skin in general. Because branched-chain amino acids are known to be required by pathogenic bacteria like *S. aureus* (*29, 30*), the ichthyotic skin could represent a more favorable environment for such bacteria. Interestingly, we also found a positive correlation between skin hydration and the branched-chain amino acids biosynthetic pathways (Fig. 5, Metacyc ID: PWY-5103, BRANCHED-CHAIN-AA-SYN-PWY, ILEUSYN-PWY) at the moist sites, suggesting that although the TGM1 genotype increased the abundance of branched-chain amino acids biosynthetic pathways, the TGM1 skin was less hydrated than the healthy skin, which dampens the enrichment. Similar to the species-level analysis, these findings showed that the ichthyosis genotype and its phenotypic manifestation can influence microbial pathways in opposing ways. Skin hydration was also positively correlated with pyruvate fermentation pathways (Metacyc ID: PWY-7111 at the moist sites and PWY-5100 at the oily sites), preQ0 biosynthetic pathway at both the oily and the moist sites (Metacyc ID: PWY-6703), and assimilatory sulfate reduction (Metacyc ID: SO4ASSIM-PWY) and L-ornithine biosynthesis pathway at the moist sites (Metacyc ID: GLUTORN-PWY) (Fig. 5 and Table S19), illustrating a complex change in the metabolic potential of the skin microbiome corresponding to changes in the skin conditions.

**Fig. 5.**
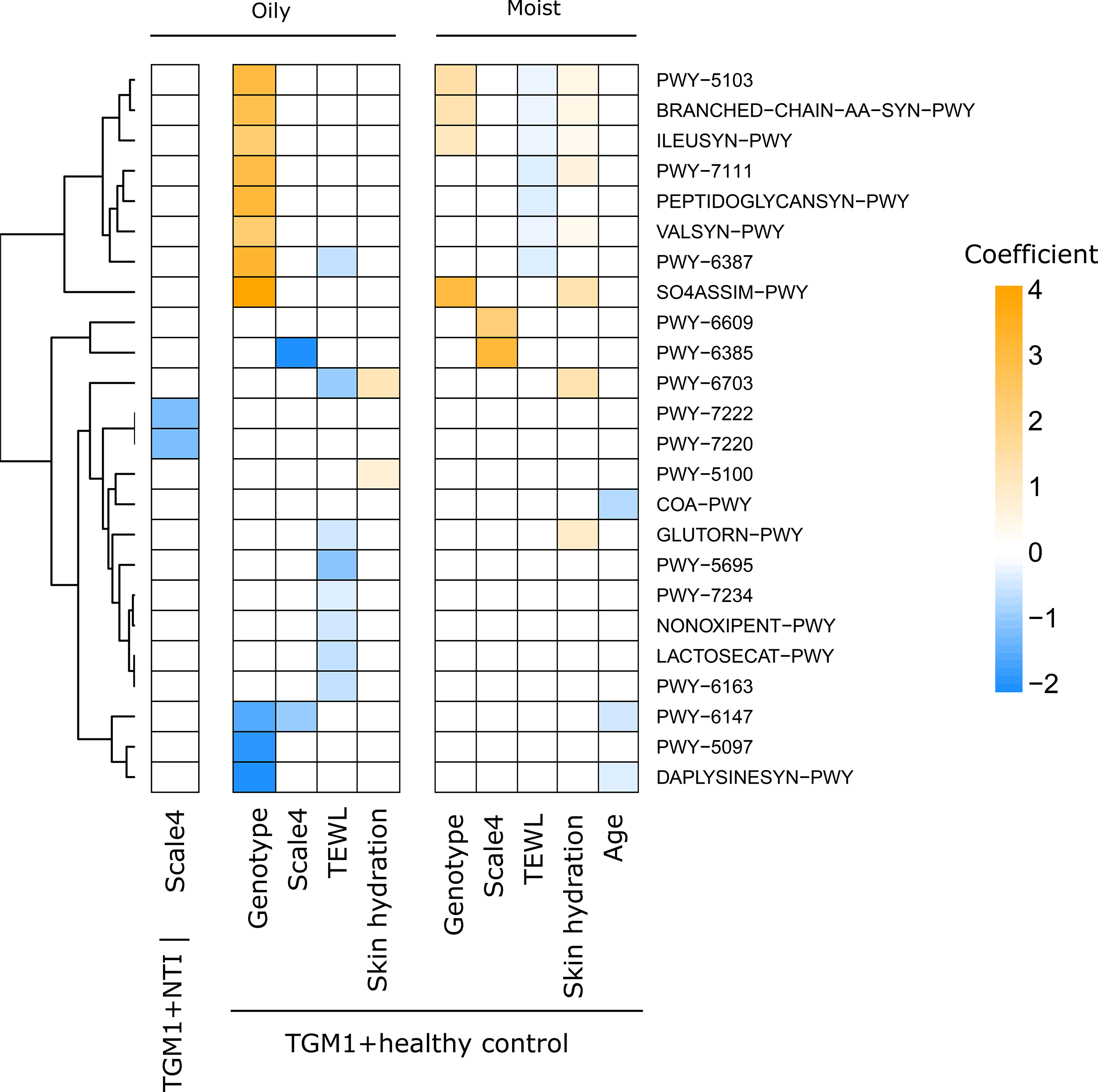
Associations between microbial functional pathways and host variables estimated using MaAsLin2. Only significant associations (q<0.25) that were concordant at the species level were shown. Values used for this figure are summarized in Table S19.

Increased TEWL, on the other hand, was negatively correlated with the abundance of multiple pathways in the TGM1+healthy control model (Fig. 5 and Table S19), demonstrating a significantly altered microbiome metabolic potential in the context of a dysfunctional stratum corneum. Many of these correlations demonstrated skin environment-specific depletion of biosynthesis pathways of compounds ranging from amino acids (Metacyc ID: PWY-5103, BRANCHED-CHAIN-AA-SYN-PWY, ILEUSYN-PWY, VALSYN-PWY) and purine (Metacyc ID: PWY-7234) to pentapeptide (Metacyc ID: PWY-6387) and peptidoglycan (Metacyc ID: PEPTIDOGLYCANSYN-PWY). The finding suggested that the nutrient acquisition strategy of skin microbes tends to transit from biosynthesis to scavenging. One pathway depleted with increased TEWL at the oily sites was the biosynthetic pathway of chorismite (Fig. 5 and Table S19, Metacyc ID: PWY-6163), a precursor of L-tryptophan, whose metabolites were known to modulate immune response via the aryl hydrocarbon receptor and plays important roles in multiple types of skin disorders (*31*). Notably, the L-tryptophan biosynthetic pathway (Metacyc ID: TRPSYN-PWY) was significantly and negatively associated with TEWL (Coefficient -0.78, q=0.022) at the oily sites in the TGM1+healthy control model, indicating that the L-tryptophan pathway abundance was altered due to the ichthyotic environment. It was not reported in Fig. 5 because the HUMAnN2 database used for the functional profiling only contained two species-level variants of the pathway (*S. epidermidis* and *S. capitis*). Although both variants were concordant (i.e. both were negatively associated with TEWL), the pathway did not meet our strict filtering criterion that a pathway needed to be represented by at least three species-level variants (detailed in the Materials and Methods section). Nonetheless, as microbial metabolites of tryptophan are involved in immune regulation in atopic dermatitis and psoriasis and have a role in skin homeostasis (*31–34*), our results suggested an additional link between tryptophan metabolism ichthyosis.

## DISCUSSION

We present here a detailed, contextualized dissection of the host genotypic and phenotypic characteristics influencing the dysbiotic ichthyosis skin microbiome, resolved at the species, strain, and metabolic pathway level. A major finding of this study is that the TGM1 genotype and the phenotypic effects associated with increased TEWL were the major drivers of microbiome diversity at the species, strain, and pathway levels. Therefore, the ichthyosis microbiome not only differs from the healthy skin microbiome due to the ichthyotic phenotype but also bears characteristics unique to the underlying genotype. Specifically, this included findings that broader skin phenotypic features such as TEWL remodeled microbiome diversity at both the species and strain-level, and the ichthyotic environment showed alterations in metabolic pathways such as branched-chain amino acids required for certain skin pathobionts and the skin barrier modulating L-tryptophan biosynthesis pathway.

One of the most substantial differences between the ichthyosis microbiome and the healthy skin microbiome is that the ichthyosis microbiome has a higher alpha diversity at the oily sites but lower alpha diversity at the moist site. At the oily sites, the difference was mainly due to the dominance of *C. acnes* in the healthy microbiome, but not in the ichthyosis microbiome, numerically decreasing the estimated alpha diversity. This observation was also reported by another group (*13*), and is likely due to the potentially less oily ichthyotic skin being unfavorable to the lipophilic *C. acnes* (*35*). However, when adjusted for covariables, Aquaphor and Vaseline treatment did not significantly increase *C. acnes* abundance, suggesting either insufficient statistical power or additional factors other than the mere abundance of lipid affecting *C. acnes*. Biologically, *C. acnes* is known to maintain skin homeostasis through immunomodulation and conferring colonization resistance (*35*), therefore its depletion could open up the skin niche to exogenous microbes. At the moist sites, however, *C. acnes* dominated neither the healthy microbiome nor the ichthyosis microbiome, and the lower diversity observed in the ichthyosis microbiome was at least partly due to stratum corneum dysfunction, as TEWL negatively correlated with diversity after adjusting for disease state. This skin environment-specific difference in microbiome diversity strongly underscores the necessity to 1) sample multiple skin niches and 2) interrogate both genotypic and phenotypic variables, which is largely missing in skin microbiome studies.

A major conclusion in our study is that skin microbiome differs not only between healthy controls and ichthyosis patients but also between different ichthyosis-causing genotypes. The difference was observed not only at the level of summary statistics, where the TGM1 microbiome tends to be the most diverse while SPINK5 tends to be the least diverse in most skin environments but also for multiple microbial species. One noteworthy example is *Co. resistens*, which was manifestly enriched in the TGM1 microbiome across almost all skin sites. Nonetheless, using multivariable regression analysis, we found that such enrichment was not driven by the TGM1 genotype alone, but was a compound effect of the TGM1 genotype and the Aquaphor treatment. On the other hand, the TGM1 genotype effect was the major driving force behind the enrichment of *S. lugdunensis*. The TGM1 genotype is characterized by a deficiency in the formation of the cornified cell envelope – an important mechanical barrier structure – surrounding corneocytes (*11, 36*), causing increased TEWL and consequently dehydration (*37*). While skin hydration in TGM1 did tend to be lower, and scale severity in TGM1 tended to be higher, than the control and other ICHT subtypes, they inadequately explain the enrichment of *S. lugdunensis* in TGM1, because the positive association was observed after correcting for skin hydration and scale severity. Therefore, *S. lugdunensis* was enriched in the TGM1 skin either because of the direct biochemical changes resulting from the TGM1 mutation, or TGM1-specific skin phenotypic changes that were not measured in this study. Although *S. lugdunensis* is known for its dual role both as part of the normal skin flora and as an opportunistic pathogen, studies of *S. lugdunensis* biology, especially in comparison to the other coagulase-negative staphylococci, are lacking, and further experimental investigations are required to mechanistically explain the genotype-microbe association.

Another important finding is that increased TEWL, the phenotypic outcome of all ichthyosis genotypes except at the sole, was a significant determinant of the structure of the ichthyosis microbiome. In this study, we found that higher TEWL – corresponding to a more severely damaged skin barrier – decreases microbiome species diversity by influencing the abundance of various bacterial and fungal species. This was consistent with the observations that skin barrier disruption was associated with dysbiotic microbiome in many skin disorders, including acne vulgaris (*38*), atopic dermatitis (*39*), and psoriasis (*40*), and our study indicated that this association extends to ichthyosis. The negative correlation between TEWL and diversity observed in acne vulgaris (*38*) was also observed in our study. Nonetheless, our study addressed the effect of skin barrier disruption as the association between TEWL and microbial taxa abundances after adjusting for the genotypic effect and host covariables, while to the best of our knowledge such deconvolution of sources of variation had not been conducted in previous studies. It is therefore unclear if TEWL in different types of diseases (or in the healthy skin) will result in the same microbiome changes. For example, we and others (*26*) observed that TEWL was positively correlated with the abundance of the *Staphylococcus* genus in the healthy control, but the individual species of *Staphylococcus* turned out to be predominantly negatively correlated with TEWL when ichthyosis samples were considered and the effect of genotype, skin hydration, scaling, and host covariables were adjusted for.

The influence of TEWL was also observed at the strain level: although the TGM1 genotype was again a major determinant of the population structure of the *Co. resistens* strains, TEWL was significantly correlated with the strain population composition of all abundant species examined in this study. However, we did not find evidence that such associations between strain abundances and phenotypes (or any other variables tested) were determined by the phylogenetic positions of the strains. Technically, our finding questions the validity of clustering phylogenetically close genomes into lineages based on the assumption that phylogenetically close genomes covary in their abundances. Biologically, this finding indicates that either 1) strain evolution in the ichthyosis environment could be highly stochastic, 2) strain evolution in the ichthyosis environment could be due to natural selection on functions whose presence were not restricted by phylogenetic positions (e.g., functions encoded by horizontally transferred genes), or 3) phylogenetically closer strains compete more often, adding a negative association to their abundances. We were unable to test these hypotheses because 1) our dataset was cross-sectional, therefore unable to assess how stochastically the strain abundances change in the ichthyosis environment, and 2) MAGs, albeit accurately reflecting the strain abundances, have limited resolution in resolving horizontally transferred genes. Future studies including time series metagenomics data or high-quality isolate genomes will be needed to test these hypotheses.

Given the possibility that ichthyotic skin shapes the microbiome through natural selection on microbial functions unrestricted by phylogenetic position, we investigated the association between genotypic variables, phenotypic variables, and host covariables with microbial functions that were present in multiple species. Interestingly, TEWL was negatively correlated with multiple biosynthetic pathways at the oily sites. The specific physiologic changes associated with increased TEWL and that favor a scavenging strategy instead of *de novo* biosynthesis are not known. Alternatively, ichthyosis can potentially alter the nutrient composition of the stratum corneum itself. These hypothetical linkage between nutrient landscape and the microbiome was well established in the nutrient-rich gut environment (*41, 42*), but not extensively studied in the nutrient-poor skin environment. Ichthyosis presents a clinically relevant scenario where altered nutrient availability potentially affects microbiome composition, which can reciprocally exacerbate the disease phenotype. This hypothesis could be experimentally investigated further using animal models or artificial skin models.

A key strength and innovation of this study is the rigorous adjustment for genotype, phenotype, and effects of variates such as treatment and age, which is seldom performed. A limitation of this study, and microbiome studies more generally, is that characterizations cannot be comprehensive, given the sequencing depth and the unclassified reads, and can be limited by sample size and data type. For example, we found changes in the abundance of *Cutibacterium* species and *M. globosa* in response to topical emollient and oral retinoid treatment to be numerically consistent with previous studies (*15, 16*), but the changes were not statistically significant. Based on our findings thus far, we suggest two additional data types that could further assist in elucidating the biological associations in the ichthyosis microbiome. First, culturomics and isolate sequencing could help explain the unclassified sequencing reads in the metagenomic samples and resolve strain-level diversity in low-abundant organisms such as *S. lugdunensis* (*20*), which was enriched in TGM1. High-quality isolate genome sequences could also reveal horizontally transferred genes that may account for the lack of phylogenetic signal in the strain-level associations identified in the present study (*43*). Second, spatial omics methods such as spatial metatranscriptomics can be useful because they effectively avoid pooling and averaging heterogenous microbiome signals at each sampled skin site (*44, 45*). This spatial resolution can be critical because 1) the ichthyosis microbiome could adapt to different regions of the scale on the diseased skin, and 2) due to the disrupted skin barrier, the ichthyosis microbiome could show higher vertical heterogeneity, which is an important mechanism in maintaining strain diversity. We also did not capture the association between ichthyosis and the abundance of fungal genera such as *Cladosporium* reported in a previous study (*19*), potentially due to the discrepancy between fungal community compositions characterized using mWGS and amplicon sequencing (*46*).

In sum, our findings revealed in high resolution how multiple aspects of ichthyosis affect the skin microbiome, especially the combined – sometimes antagonistic – influence of the ichthyosis genotypes and the increased TEWL phenotype. We also generated a testable hypothesis that ichthyosis in general shapes the skin microbiome through an altered nutrient landscape, findings only possible by simultaneously adjusting for the effects of topical/oral treatments and host age on the skin microbes.

## MATERIALS AND METHODS

### Study design

36 patients with ichthyosis and 14 controls were enrolled. All controls and 18 ichthyosis patients were sampled at the Yale Dermatology clinic; 18 additional ichthyosis patients were sampled at meetings of the Foundation for Ichthyosis and Related Skin Types. All subjects were at least 18 years of age and had used no oral or topical antibiotic in the month prior to sampling; they used only Dove soap for bathing for one week prior to sampling; they applied no emollients for 24 hours prior to sampling. Ichthyosis subjects had genetic sequence-confirmed causative mutations in either KRT1, KRT2, KRT10, TGM1, ABCA12, ALOX12B, NIPAL4, PNPLA1, or SPINK5. Except for patients with TGM1 mutations, only two patients (ICHT-1 and ICHT-38) with different disease genotypes were taking oral retinoids. They were excluded from the analysis to avoid a skewed representation of the effect of retinoids. All subjects gave informed consent for this Yale IRB-approved study (HIC #2000023381). Swabs for microbial DNA were obtained from 10 body sites (scalp vertex; post-auricular scalp; forehead; upper lateral arm; upper back; central chest; sacrum above gluteal cleft; popliteal fossa; shin; sole) and an air control using established procedures (*20*). Each site was evaluated for the degree of scale as previously described (*47*). Measurements of skin hydration and TEWL were obtained at each site prior to swabbing using Courage+Khazaka electronic probes.

### mWGS sequencing and preprocessing

DNA was extracted with a bead-beating + enzymatic lysis manual extraction method and sequencing libraries were prepared using ¼ volume reactions of the Illumina DNA Prep kit and Nextera Unique Dual Indexes (Illumina), as previously described (*20*). Pooled libraries were sequenced on a NovaSeq 6000 (Illumina). Demultiplexed Illumina reads were trimmed with Trimmomatic (v0.39) (*48*), with parameters: seed mismatch=2, palindrome clip threshold=30, simple clip threshold=10, and LEADING:3, TRAILING:3, SLIDINGWINDOW:4:15, MINLEN:36. Human reads were then removed with Bowtie2 (v2.2.9, -<u>-</u>very-sensitive mode) (*49*), mapping to the CHM13 (v2.0) human reference genome (*50*).

### Assembly and quality filtering of MAGs

Metagenomic samples were first assembled *de novo* using MEGAHIT (v1.0.6) (*51, 52*) with default parameters. The resulting assemblies were filtered to only include contigs no shorter than 1000 base pairs. Metagenomic reads were then mapped back to the contigs using Bowtie2 (v2.3.4.3) (*49*) under “very-sensitive” mode. The resulting SAM files were binarized and sorted using SAMtools (v1.10) (*53*) before used for binning through MetaBAT2 (v2.12.1) (*54*) with the option “-m 1500”. The resulting raw genome bins (N=5906) were quality-filtered using checkM (v1.1.2) (*55*) to keep bins of at least medium quality (genome completeness no lower than 50% and contamination no higher than 10%). These bins (N=2136) were pairwise compared using dRep (v2.4.0) (*56*) to identify and remove duplicated bins (average nucleotide identity greater than 99% and level of overlap greater than 75%). We prioritized removing those duplicated bins that had lower genome completeness. When duplicates had the same completeness, we removed the ones with the higher contamination. The resulting 907 non-redundant bins of at least medium quality were referred to as MAGs (metagenomic assembled genomes). Taxonomic annotations and phylogenetic relationships of the MAGs were estimated using the classify workflow of GTDB-TK (v1.6.0) (*57*).

### Taxonomic profiling

Species-level community compositions were profiled using MetaPhlAn4.0 (*24*). To estimate strain-level compositions, we used a reference-based method modified from Larson et al. (2022). A reference database was generated by compiling all MAGs assembled from the dataset (detailed below). Sequences for these MAGs were available at https://github.com/ohlab/ICHT_MAGs. Reads were then mapped to the reference database using Bowtie2 (v2.3.4.3) (*49*) with k=10 and “very-sensitive” mode. We then used Pathoscope (v2.0.6) (*58*) on the resulting SAM files using default parameters to reassign reads to the most likely genome of origin.

### Metabolic pathway profiling

To characterize the microbial metabolic potentials in our dataset, reads were processed using HUMAnN2 (v2.8.0, diamond mode) (*59*) to compute the abundances of microbial metabolic pathways. The resulting abundance matrixes were then used for multivariable regression analyses using MaAsLin2 (v1.8.0, detailed below) (*27*). To identify pathway associations that were more likely functionally relevant, instead of merely “hitchhiking” with other fitness-changing genes, we filtered our results to focus on pathways that were 1) significant at the community level, as defined by HUMAnN2, and 2) concordant at the species level: for a given association between a community-level pathway and a host variable, the pathway needs to be represented by at least 3 species-level variants, and at least 75% of the species-level variants need to be correlated with the variable in the same direction as the community-level pathway.

### Statistical analysis

All statistical analyses were performed in R (4.1.3) (*60*) unless otherwise noted. False discovery rates were controlled using the Benjamini-Hochberg procedure. We consider statistical tests with p (or q) < 0.05 to be significant, except when running MaAsLin2 (*27*), where the default threshold of q < 0.25 was adopted. Wilcoxon Rank Sum test was conducted using the “wilcox.test” function in R. Cophenetic distances were computed using the “cophenetic.phylo” function in the R package “ape” (v5.6) (*61*). Alpha diversity was projected using Shannon’s diversity index via the “diversity” function in the R package “vegan” (v2.5.7) (*62*). We used mixed effect model (the lmer function in the R package “lme4” (v1.1) (*63*) to test the association between host variables and alpha diversity. To respect the biogeographic diversity associated with different skin environments, we fit a separate mixed effect model for each skin type (dry, oily, moist, or sole). In each model, ichthyosis genotype (TGM1 or healthy controls/NTI), skin hydration (corneometer measurement), TEWL (tewameter measurement), whether scale severity is ≥ 2, whether scale severity is ≥ 3, whether scale severity is ≥ 4, Vaseline treatment, Aquaphor treatment, Retinoids treatment, and age were modeled as fixed effects. In dry, oily, and moist models, subject and skin sites were modeled as random effects to account for the greater variation between subjects/sampling locations than within a subject/sampling location. Because of its ordinality, scale severity was represented using three binary variables instead of one numerical variable. Significance of the predictors was tested using type II ANOVA tests using the “Anova” function in the R package “lmerTest” (v3.1.3) (*64*). Nonparametric bootstrap was implemented using the “two.boot” function in the R package simpleboot (v1.1) (*65*) with 1000 bootstrap replications for each mean comparison. The confidence intervals were then estimated using the “boot.ci” function in the R package boot (v1.3) (Davison and Hinkley 1997; Canty and support 2022). Multivariable regression models were fit using MaAsLin2 (v1.8.0) (*27*), with options standardize=T, min_abundance=0.1 (corresponding to relative abundance no lower than 0.1%) for species abundance profiles, min_abundance=0 for MAG abundance profiles, and min_abundance=1 (corresponding to RPKM no lower than 1). To accounting for biogeographic heterogeneity, we fit a separate model for each skin type (dry, oily, and moist). Like described above, in each model, ichthyosis genotype (TGM1 or healthy controls/NTI), skin hydration (corneometer measurement), TEWL (tewameter measurement), whether scale severity is ≥ 2, whether scale severity is ≥ 3, whether scale severity is ≥ 4, Vaseline treatment, Aquaphor treatment, Retinoids treatment, and age were modeled as fixed effects. In dry, oily, and moist models, subject and skin sites were modeled as random effects to account for the greater variation between subjects/sampling locations than within a subject/sampling location. Because of its ordinality, scale severity was represented using three binary variables instead of one numerical variable.

## Acknowledgments

We are grateful to all patients and healthy individuals that participated in the study. We wish to thank Dr. Erin Mathes and Dr. Evelyn Lilly for referring patients.

## Funding

National Institutes of Health grant 1 DP2 GM126893-01 (JO) National Institutes of Health grant 5 R21 AR075174 (JO)

## Author contributions

Conceptualization: JO, LM

Methodology: JO, LM, WZ, NR

Investigation: RC, WZ, LM, NR

Visualization: WZ

Funding acquisition: JO, LM

Project administration: JO, LM Supervision: JO, LM

Writing – original draft: WZ

Writing – review & editing: JO, LM, WZ

## Competing interests

Authors declare that they have no competing interests.

## Data and materials availability

mWGS data (trimmed) can be accessed in National Center for Biotechnology Information (NCBI) BioProject PRJNA1045259. MAGs can be accessed from https://github.com/ohlab/icht_MAGs. Computational code for data analyses is available upon reasonable request.

## Figures

**Fig. S1.**
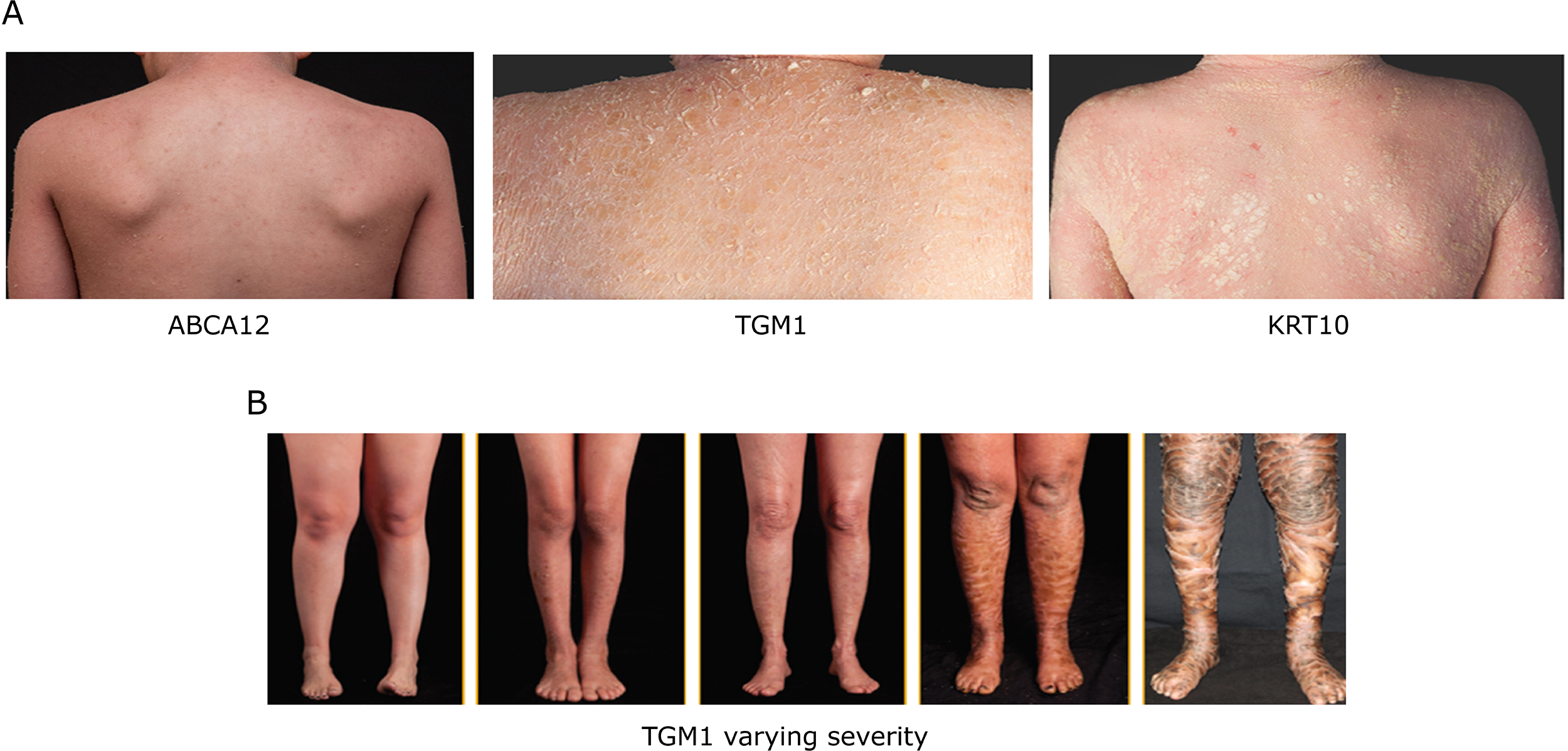
Severity and quality of scale vary in ichthyosis. (**A**) Skin with an *ABCA12* mutation usually has small flaky scales with background erythema. This phenotype was formally known as congenital ichthyosiform erythroderma. Skin with a *TGM1* mutation usually has large, plate-like scales and mild erythema. This phenotype was formerly known as lamellar ichthyosis. Skin with a *KRT10* mutation usually has cobble-stone-like scale in a corrugated pattern and mild erythema. This phenotype is characteristic of epidermolytic ichthyosis. (**B**) Shins of five patients with *TGM1* mutation demonstrating the varying severity scale.

**Fig. S2.**
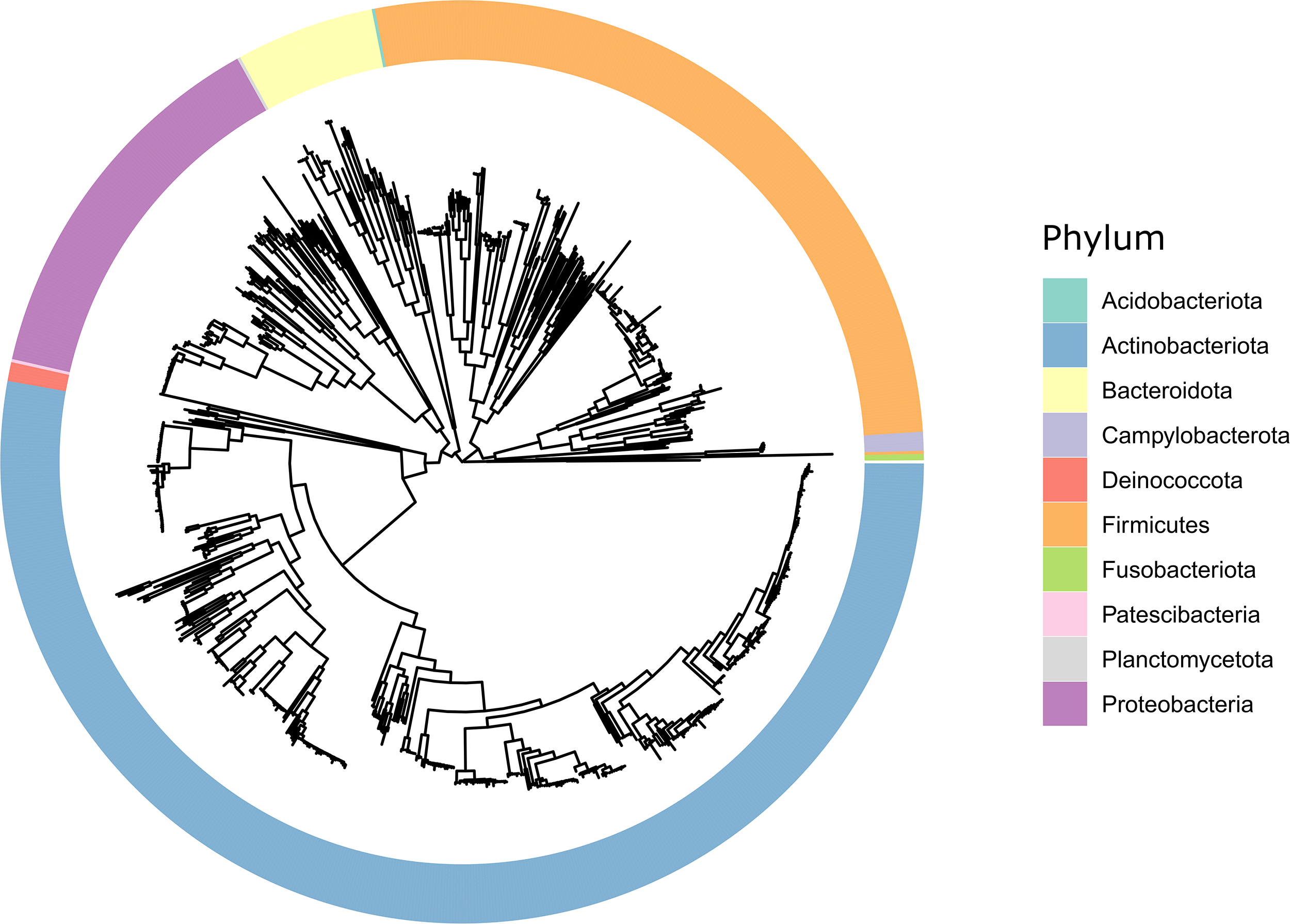
phylogenetic relationships of the 907 MAGs used for the analysis.

**Supplementary Tables:**

https://www.dropbox.com/scl/fi/fys2urkmai0kghis8lob7/SupplementaryTables.xlsx?rlkey=ciopn4fm0wx8d3atcwjb1g3id&dl=0

## Supplementary Note

Here we investigated skin microbiome signatures at the species, strain, and metabolic pathway levels in five ichthyosis genotypes (KRT2, KRT10, ABCA12, ALOX12B, and NIPAL4). Due to the rarity of these genotypes, we based the bulk of our analyses on grouping the genotypes by their direct effect on critical components of the stratum corneum: keratinopathic ichthyoses (KPI) affect the production of keratin filaments and are represented by KRT2 and KRT10, and lipid function ichthyoses (LFI) affect lipid metabolism/transfer functions and are represented by ABCA12, ALOX12B, and NIPAL4.

We first sought to identify microbial species manifestly associated with the control, TGM1, KPI, or LFI with at least a 95% confidence using nonparametric bootstrap. Our findings were consistent with what was observed when each genotype was analyzed separately (Supplementary Fig. 1A and Fig. 1C): *Cutibacterium (C.) acnes* and *Malassezia (M.) globosa* were enriched in the healthy controls; *Corynebacterium (Co.) resistens, Staphylococcus (S.) lugdunensis, Prevotella (P.) disiens*, and *Co. bovis* associated with TGM1; microbe associations with KPI and LFI appeared more nuanced and skin site-specific.

We then investigated which genotypic group, phenotype, and host covariables are associated with each microbial species constituting the skin microbiome, using the multivariable regression method implemented in MaAsLin2. We found that skin species were influenced by genotypic group, phenotype, and host covariables collectively (Supplementary Note Figure 1B and Supplementary Note Table 1). For example, the relative abundance of *S. haemolyticus* was higher in LFI but lowered when treated with Vaseline (Supplementary Note Figure 1B and Supplementary Note Table 1). We found few skin species significantly influenced by the KPI genotypes, but Aquaphor treatment in KPI ichthyosis patients perturbed the relative abundance of multiple species: *Co. bovis*, *Co. massiliense*, and *S. capitis* exhibited elevated relative abundances with Aquaphor treatments in multiple skin types; while *S. hominis*, *S. epidermidis*, *C. granulosum*, and *C. acnes* showed significantly decreased relative abundance with Aquaphor treatments in multiple skin types (Supplementary Note Figure 1A and Supplementary Note Table 1). Finally, both TEWL and skin hydration significantly impacted multiple species in both genotypic groups, while these impacts were often skin environment-specific (Supplementary Note Figure 1B and Supplementary Note Table 1).

We next asked if genotypic group, phenotype, and host covariables influence the structure of microbial strain populations in LFI and KPI patients. Similar to the strain-level analysis on TGM1 samples, we focused on MAGs from four species (*C. acnes*, *Co. resistens*, *Co. amycolatum*, and *S. capitis*) that had most of that species’ strain variation represented by the MAGs. Consistent with the TGM1 analysis, we found few significant associations between MAG abundances and genotypic group or host covariables, while the phenotypic variable TEWL negatively correlated with multiple MAGs in all four species (Supplementary Note Figure 1C and Supplementary Note Table 2-5). This result supports the hypothesis that skin barrier disruption can cause strain-level perturbations across ichthyosis genotypes. Also consistent with the findings from TGM1 patients was that, for LFI and KPI patients, 1) the phylogenetic similarity of MAGs was in general not positively correlated with the MAGs’ associations with genotype, phenotype, and host covariables (Supplementary Note Figure 1D), and 2) the correlation coefficients between phylogenetic differences and the differences in associations with host variables tended to be slightly negative for *S. capitis* MAGs (Supplementary Note Figure 1D). This again brought about the possibility that phylogenetically close *S. capitis* strains could interact more antagonistically than phylogenetically distant strains.

We next characterized microbial metabolic pathways that were associated with KPI or LFI, phenotype, and host covariables. Similar to the criterion we used for the TGM1 samples, we focused on microbial metabolic pathway associations that were not only significant on the community level but also concordant at the species level. With the strict criterion, we were only able to identify associations at the moist and oily sites. We found more genotype-, phenotype- and covariable-associated pathways in the LFI+healthy controls model than the KPI+healthy controls model (Supplementary Note Figure 1E and Supplementary Note Table 6). These included biosynthetic pathways of many amino acids – L-isoleucine (MetaCyc ID: PWY-5103, ILEUSYN-PWY, and PWY-3001), L-arginine (MetaCyc ID: ARGSYN-PWY), L-proline (MetaCyc ID: PWY-4981), and branched-chain amino acids (MetaCyc ID: BRANCHED-CHAIN-AA-SYN-PWY) (Supplementary Note Figure 1E and Supplementary Note Table 6). Consistent with the findings from the TGM1 samples, these amino acids biosynthetic pathways were often positively associated with the genotype effect of LFI but negatively associated with TEWL (Supplementary Note Figure 1E and Supplementary Note Table 6), suggesting that skin barrier disruption could be in favor of scavenging rather than biosynthesis of amino acids. Also consistent with the findings from the TGM1 samples was that TEWL was negatively associated with chorismite biosynthesis pathway (MetaCyc ID: PWY-6163) at the oily sites in both the KPI and LFI samples (Supplementary Note Figure 1E and Supplementary Note Table 6). As chorismite was a precursor of L-tryptophan, this association suggested that tryptophan metabolism could be related to barrier disruption caused by ichthyosis.

## Supplementary Note Figures

**Supplementary Note Fig. 1.**
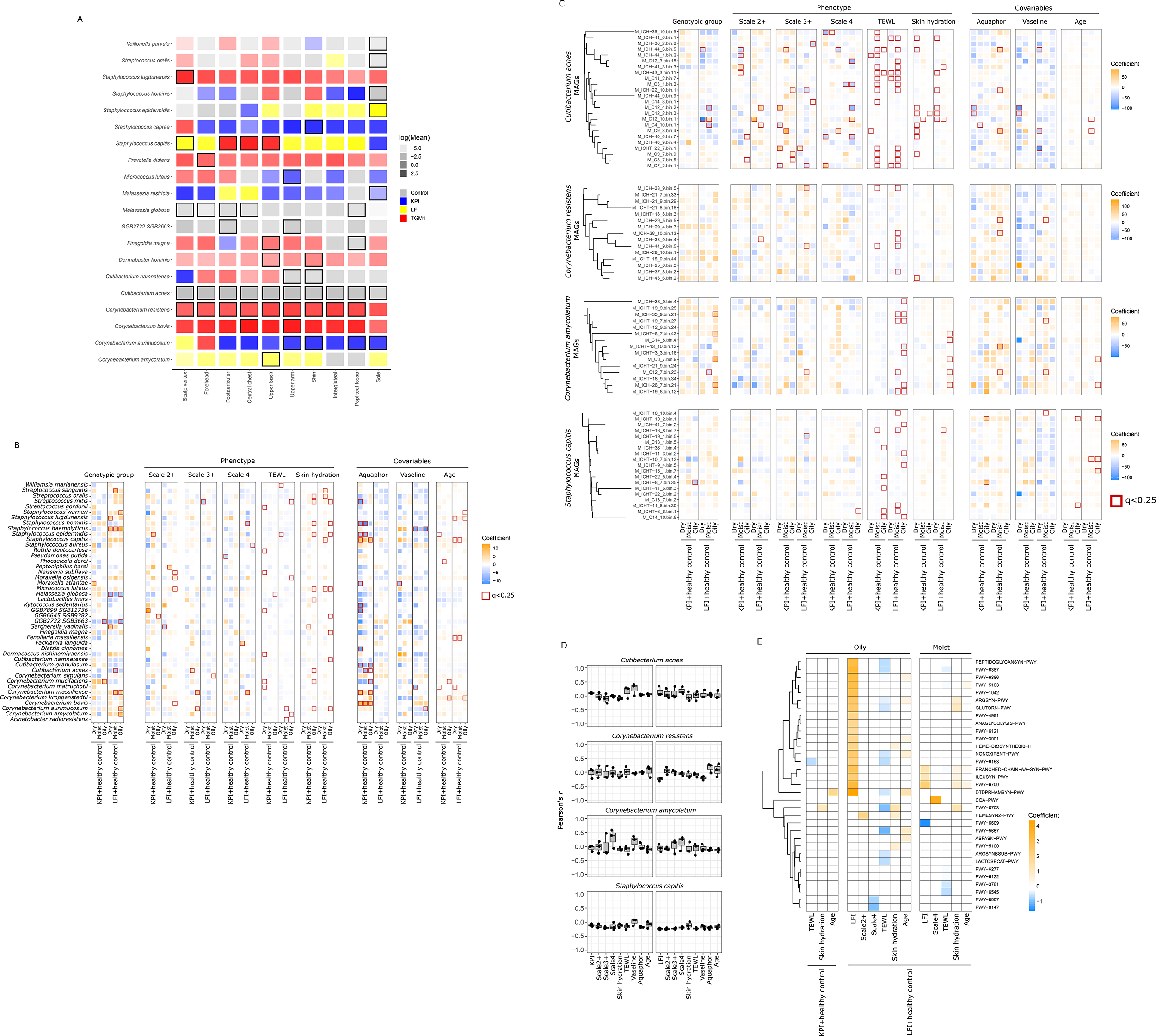
Skin microbiome signatures in LFI and KPI. (A) Manifest enrichment of microbial species in the control, KPI, LFI, or TGM1 microbiome. Transparency of the tile colors represents the mean difference in species relative abundance in one group versus all other samples. Tiles with black outlines represent species where the lower bound of the 95% confidence interval of the mean difference was greater than zero. (B) Associations between species relative abundances and host variables estimated using MaAsLin2. Significant associations (q < 0.25) had red outlines. Only species with at least one significant association were shown. Values used for this figure were summarized in Supplementary Note Table 1. (C) Associations between the relative abundances of MAGs from four abundant species and host variables estimated using MaAsLin2. The relative abundances of the MAGs were rescaled within each species to sum up to 1. Phylogenetic relationships of the MAGs from each of the four species were illustrated using a subtree of Figure S2. Significant associations (q < 0.25) had red outlines. Values used for this figure were summarized in Supplementary Note Table 2-5. (D) Pearson’s correlation coefficient between cophenetic distance of MAGs and the MAGs’ differential response to host variables. Each data point represents a skin environment (dry, oily, or moist). (E) Associations between microbial functional pathways and host variables estimated using MaAsLin2. Only significant associations (q<0.25) that were concordant at the species level were shown. Values used for this figure were summarized in Supplementary Note Table 6.

**Supplementary Note Tables:**

https://www.dropbox.com/scl/fi/rfd09pb747tdavwyito6g/SupplementaryNoteTables.xlsx?rlkey=io04t8ayw1zgzyzlglz9r999l&dl=0

## Notes

### Competing Interest Statement

The authors have declared no competing interest.

## Reference

1. H.-J. Lee, M. Kim, Skin Barrier Function and the Microbiome. Int J Mol Sci 23, 13071 (2022).

2. N. Lunjani, S. Ahearn-Ford, F. S. Dube, C. Hlela, L. O’Mahony, Mechanisms of microbe-immune system dialogue within the skin. Genes Immun 22, 276–288 (2021).

3. Q. Liu, R. Ranallo, C. Rios, E. A. Grice, K. Moon, R. L. Gallo, Crosstalk between skin microbiota and immune system in health and disease. Nat Immunol 24, 895–898 (2023).

4. A. L. Cogen, V. Nizet, R. L. Gallo, Skin microbiota: a source of disease or defence? The British journal of dermatology 158, 442 (2008).

5. K. Findley, J. Oh, J. Yang, S. Conlan, C. Deming, J. A. Meyer, D. Schoenfeld, E. Nomicos, M. Park, NIH Intramural Sequencing Center Comparative Sequencing Program, H. H. Kong, J. A. Segre, Topographic diversity of fungal and bacterial communities in human skin. Nature 498, 367–370 (2013).

6. J. Oh, A. L. Byrd, C. Deming, S. Conlan, NISC Comparative Sequencing Program, H. H. Kong, J. A. Segre, Biogeography and individuality shape function in the human skin metagenome. Nature 514, 59–64 (2014).

7. R. Blekhman, J. K. Goodrich, K. Huang, Q. Sun, R. Bukowski, J. T. Bell, T. D. Spector, A. Keinan, R. E. Ley, D. Gevers, A. G. Clark, Host genetic variation impacts microbiome composition across human body sites. Genome Biology 16, 191 (2015).

8. L. Moitinho-Silva, F. Degenhardt, E. Rodriguez, H. Emmert, S. Juzenas, L. Möbus, F. Uellendahl-Werth, N. Sander, H. Baurecht, L. Tittmann, W. Lieb, C. Gieger, A. Peters, D. Ellinghaus, C. Bang, A. Franke, S. Weidinger, M. C. Rühlemann, Host genetic factors related to innate immunity, environmental sensing and cellular functions are associated with human skin microbiota. Nat Commun 13, 6204 (2022).

9. C. Gutiérrez-Cerrajero, E. Sprecher, A. S. Paller, M. Akiyama, J. Mazereeuw-Hautier, A. Hernández-Martín, R. González-Sarmiento, Ichthyosis. Nat Rev Dis Primers 9, 2 (2023).

10. K. A. Choate, L. M. Milstone, in Fitzpatrick’s Dermatology, S. Kang, M. Amagai, A. L. Bruckner, A. H. Enk, D. J. Margolis, A. J. McMichael, J. S. Orringer, Eds. (McGraw-Hill Education, New York, NY, 2019).

11. Q. Sun, N. M. Burgren, S. Cheraghlou, A. S. Paller, M. Larralde, L. Bercovitch, J. Levinsohn, I. Ren, R. H. Hu, J. Zhou, T. Zaki, R. Fan, C. Tian, C. Saraceni, C. J. Nelson-Williams, E. Loring, B. G. Craiglow, L. M. Milstone, R. P. Lifton, L. M. Boyden, K. A. Choate, The Genomic and Phenotypic Landscape of Ichthyosis: An Analysis of 1000 Kindreds. JAMA Dermatol 158, 16–25 (2022).

12. P. L. J. M. Zeeuwen, T. H. A. Ederveen, D. A. van der Krieken, H. Niehues, J. Boekhorst, S. Kezic, D. A. T. Hanssen, M. E. Otero, I. M. J. J. van Vlijmen-Willems, D. Rodijk-Olthuis, D. Falcone, E. H. J. van den Bogaard, M. Kamsteeg, H. D. de Koning, M. E. J. Zeeuwen-Franssen, M. A. M. van Steensel, M. Kleerebezem, H. M. Timmerman, S. A. F. T. van Hijum, J. Schalkwijk, Gram-positive anaerobe cocci are underrepresented in the microbiome of filaggrin-deficient human skin. Journal of Allergy and Clinical Immunology 139, 1368–1371 (2017).

13. K.-C. Tham, R. Lefferdink, K. Duan, S. S. Lim, X. F. C. C. Wong, E. Ibler, B. Wu, H. Abu-Zayed, S. M. Rangel, E. Del Duca, M. Chowdhury, M. Chima, H. J. Kim, B. Lee, E. Guttman-Yassky, A. S. Paller, J. E. A. Common, Distinct skin microbiome community structures in congenital ichthyosis. Br J Dermatol 187, 557–570 (2022).

14. W. H. McCoy, E. Otchere, B. A. Rosa, J. Martin, C. M. Mann, M. Mitreva, Skin Ecology during Sebaceous Drought-How Skin Microbes Respond to Isotretinoin. J Invest Dermatol 139, 732–735 (2019).

15. H.-L. Kelhälä, V. T. E. Aho, N. Fyhrquist, P. A. B. Pereira, M. E. Kubin, L. Paulin, R. Palatsi, P. Auvinen, K. Tasanen, A. Lauerma, Isotretinoin and lymecycline treatments modify the skin microbiota in acne. Exp Dermatol 27, 30–36 (2018).

16. S. Seite, G. E. Flores, J. B. Henley, R. Martin, H. Zelenkova, L. Aguilar, N. Fierer, Microbiome of affected and unaffected skin of patients with atopic dermatitis before and after emollient treatment. J Drugs Dermatol 13, 1365–1372 (2014).

17. P. J. Larson, W. Zhou, A. Santiago, S. Driscoll, E. Fleming, A. Y. Voigt, O. K. Chun, J. J. Grady, G. A. Kuchel, J. T. Robison, J. Oh, Associations of the skin, oral and gut microbiome with aging, frailty and infection risk reservoirs in older adults. Nat Aging 2, 941–955 (2022).

18. W. Zhou, E. Fleming, G. Legendre, L. Roux, J. Latreille, G. Gendronneau, S. Forestier, J. Oh, Skin microbiome attributes associate with biophysical skin ageing. Exp Dermatol 32, 1546–1556 (2023).

19. V. Moosbrugger-Martinz, H. Hackl, R. Gruber, M. Pilecky, L. Knabl, D. Orth-Höller, S. Dubrac, Initial Evidence of Distinguishable Bacterial and Fungal Dysbiosis in the Skin of Patients with Atopic Dermatitis or Netherton Syndrome. J Invest Dermatol 141, 114–123 (2021).

20. W. Zhou, M. Spoto, R. Hardy, C. Guan, E. Fleming, P. J. Larson, J. S. Brown, J. Oh, Host-specific evolutionary and transmission dynamics shape the functional diversification of Staphylococcus epidermidis in human skin. Cell 180, 454–470.e18 (2020).

21. R. Caldwell, W. Zhou, J. Oh, Strains to go: interactions of the skin microbiome beyond its species. Curr Opin Microbiol 70, 102222 (2022).

22. K. Ide, T. Saeki, K. Arikawa, T. Yoda, T. Endoh, A. Matsuhashi, H. Takeyama, M. Hosokawa, Exploring strain diversity of dominant human skin bacterial species using single-cell genome sequencing. Frontiers in Microbiology 13 (2022) (available at https://www.frontiersin.org/articles/10.3389/fmicb.2022.955404).

23. A. Conwill, A. C. Kuan, R. Damerla, A. J. Poret, J. S. Baker, A. D. Tripp, E. J. Alm, T. D. Lieberman, Anatomy promotes neutral coexistence of strains in the human skin microbiome. Cell Host Microbe 30, 171–182.e7 (2022).

24. A. Blanco-Míguez, F. Beghini, F. Cumbo, L. J. McIver, K. N. Thompson, M. Zolfo, P. Manghi, L. Dubois, K. D. Huang, A. M. Thomas, W. A. Nickols, G. Piccinno, E. Piperni, M. Punčochář, M. Valles-Colomer, A. Tett, F. Giordano, R. Davies, J. Wolf, S. E. Berry, T. D. Spector, E. A. Franzosa, E. Pasolli, F. Asnicar, C. Huttenhower, N. Segata, Extending and improving metagenomic taxonomic profiling with uncharacterized species using MetaPhlAn 4. Nat Biotechnol, 1–12 (2023).

25. T. R. Erickson, M. B. Murphrey, H. Abu-Zayed, B. Wu, E. Ibler, S. M. Rangel, A. S. Paller, Transepidermal water loss in the orphan forms of ichthyosis. Pediatric Dermatology 37, 771–773 (2020).

26. J.-H. Kim, S.-M. Son, H. Park, B. K. Kim, I. S. Choi, H. Kim, C. S. Huh, Taxonomic profiling of skin microbiome and correlation with clinical skin parameters in healthy Koreans. Sci Rep 11, 16269 (2021).

27. H. Mallick, A. Rahnavard, L. J. McIver, S. Ma, Y. Zhang, L. H. Nguyen, T. L. Tickle, G. Weingart, B. Ren, E. H. Schwager, S. Chatterjee, K. N. Thompson, J. E. Wilkinson, A. Subramanian, Y. Lu, L. Waldron, J. N. Paulson, E. A. Franzosa, H. C. Bravo, C. Huttenhower, Multivariable association discovery in population-scale meta-omics studies. PLoS Comput Biol 17, e1009442 (2021).

28. A. L. Byrd, C. Deming, S. K. B. Cassidy, O. J. Harrison, W.-I. Ng, S. Conlan, Y. Belkaid, J. A. Segre, H. H. Kong, Staphylococcus aureus and S. epidermidis strain diversity underlying human atopic dermatitis. Sci Transl Med 9, eaal4651 (2017).

29. H. Chen, Q. Zhao, Q. Zhong, C. Duan, J. Krutmann, J. Wang, J. Xia, Skin Microbiome, Metabolome and Skin Phenome, from the Perspectives of Skin as an Ecosystem. Phenomics 2, 363–382 (2022).

30. J. C. Kaiser, A. N. King, J. C. Grigg, J. R. Sheldon, D. R. Edgell, M. E. P. Murphy, S. R. Brinsmade, D. E. Heinrichs, Repression of branched-chain amino acid synthesis in Staphylococcus aureus is mediated by isoleucine via CodY, and by a leucine-rich attenuator peptide. PLoS Genet 14, e1007159 (2018).

31. V. Jiminez, N. Yusuf, Bacterial Metabolites and Inflammatory Skin Diseases. Metabolites 13, 952 (2023).

32. A. Uberoi, C. Bartow-McKenney, Q. Zheng, L. Flowers, A. Campbell, S. A. B. Knight, N. Chan, M. Wei, V. Lovins, J. Bugayev, J. Horwinski, C. Bradley, J. Meyer, D. Crumrine, C. H. Sutter, P. Elias, E. Mauldin, T. R. Sutter, E. A. Grice, Commensal microbiota regulates skin barrier function and repair via signaling through the aryl hydrocarbon receptor. Cell Host & Microbe 29, 1235–1248.e8 (2021).

33. N. Fernández-Gallego, F. Sánchez-Madrid, D. Cibrian, Role of AHR Ligands in Skin Homeostasis and Cutaneous Inflammation. Cells 10, 3176 (2021).

34. M. Szelest, K. Walczak, T. Plech, A New Insight into the Potential Role of Tryptophan-Derived AhR Ligands in Skin Physiological and Pathological Processes. Int J Mol Sci 22, 1104 (2021).

35. C. M. Ahle, C. Feidenhansl, H. Brüggemann, Cutibacterium acnes. Trends in Microbiology 31, 419– 420 (2023).

36. M. L. Herman, S. Farasat, P. J. Steinbach, M.-H. Wei, O. Toure, P. Fleckman, P. Blake, S. J. Bale, J. R. Toro, Transglutaminase-1 (TGM1) Gene Mutations in Autosomal Recessive Congenital Ichthyosis: Summary of Mutations (Including 23 Novel) and Modeling of TGase-1. Hum Mutat 30, 537–547 (2009).

37. K. Aufenvenne, F. Larcher, I. Hausser, B. Duarte, V. Oji, H. Nikolenko, M. Del Rio, M. Dathe, H. Traupe, Topical Enzyme-Replacement Therapy Restores Transglutaminase 1 Activity and Corrects Architecture of Transglutaminase-1-Deficient Skin Grafts. Am J Hum Genet 93, 620–630 (2013).

38. L. Zhou, X. Liu, X. Li, X. He, X. Xiong, J. Lai, Epidermal Barrier Integrity is Associated with Both Skin Microbiome Diversity and Composition in Patients with Acne Vulgaris. Clin Cosmet Investig Dermatol 15, 2065–2075 (2022).

39. C. L. Jinnestål, E. Belfrage, O. Bäck, A. Schmidtchen, A. Sonesson, Skin barrier impairment correlates with cutaneous Staphylococcus aureus colonization and sensitization to skin-associated microbial antigens in adult patients with atopic dermatitis. Int J Dermatol 53, 27–33 (2014).

40. W.-M. Wang, H.-Z. Jin, Skin Microbiome: An Actor in the Pathogenesis of Psoriasis. Chin Med J (Engl) 131, 95–98 (2018).

41. F. C. Pereira, D. Berry, Microbial nutrient niches in the gut. Environ Microbiol 19, 1366–1378 (2017).

42. P. Zhang, Influence of Foods and Nutrition on the Gut Microbiome and Implications for Intestinal Health. Int J Mol Sci 23, 9588 (2022).

43. M. Groussin, M. Poyet, A. Sistiaga, S. M. Kearney, K. Moniz, M. Noel, J. Hooker, S. M. Gibbons, L. Segurel, A. Froment, R. S. Mohamed, A. Fezeu, V. A. Juimo, S. Lafosse, F. E. Tabe, C. Girard, D. Iqaluk, L. T. T. Nguyen, B. J. Shapiro, J. Lehtimäki, L. Ruokolainen, P. P. Kettunen, T. Vatanen, S. Sigwazi, A. Mabulla, M. Domínguez-Rodrigo, Y. A. Nartey, A. Agyei-Nkansah, A. Duah, Y. A. Awuku, K. A. Valles, S. O. Asibey, M. Y. Afihene, L. R. Roberts, A. Plymoth, C. A. Onyekwere, R. E. Summons, R. J. Xavier, E. J. Alm, Elevated rates of horizontal gene transfer in the industrialized human microbiome. Cell 184, 2053–2067.e18 (2021).

44. L. Lyu, X. Li, R. Feng, X. Zhou, T. K. Guha, X. Yu, G. Q. Chen, Y. Yao, B. Su, D. Zou, M. P. Snyder, L. Chen, Simultaneous profiling of host expression and microbial abundance by spatial metatranscriptome sequencing. Genome Res. 33, 401–411 (2023).

45. A. Wong-Rolle, Q. Dong, Y. Zhu, P. Divakar, J. L. Hor, N. Kedei, M. Wong, D. Tillo, E. A. Conner, Rajan, D. S. Schrump, C. Jin, R. N. Germain, C. Zhao, Spatial meta-transcriptomics reveal associations of intratumor bacteria burden with lung cancer cells showing a distinct oncogenic signature. J Immunother Cancer 10, e004698 (2022).

46. M. Usyk, B. A. Peters, S. Karthikeyan, D. McDonald, C. C. Sollecito, Y. Vazquez-Baeza, J. P. Shaffer, M. D. Gellman, G. A. Talavera, M. L. Daviglus, B. Thyagarajan, R. Knight, Q. Qi, R. Kaplan, R. D. Burk, Comprehensive evaluation of shotgun metagenomics, amplicon sequencing, and harmonization of these platforms for epidemiological studies. Cell Rep Methods 3, 100391 (2023).

47. N. V. Marukian, Y. Deng, G. Gan, I. Ren, W. Thermidor, B. G. Craiglow, L. M. Milstone, K. A. Choate, Establishing and Validating an Ichthyosis Severity Index. J Invest Dermatol 137, 1834–1841 (2017).

48. A. M. Bolger, M. Lohse, B. Usadel, Trimmomatic: a flexible trimmer for Illumina sequence data. Bioinformatics 30, 2114–2120 (2014).

49. B. Langmead, S. L. Salzberg, Fast gapped-read alignment with Bowtie 2. Nat Methods 9, 357–359 (2012).

50. S. Nurk, S. Koren, A. Rhie, M. Rautiainen, A. V. Bzikadze, A. Mikheenko, M. R. Vollger, N. Altemose, L. Uralsky, A. Gershman, S. Aganezov, S. J. Hoyt, M. Diekhans, G. A. Logsdon, M. Alonge, S. E. Antonarakis, M. Borchers, G. G. Bouffard, S. Y. Brooks, G. V. Caldas, N.-C. Chen, H. Cheng, C.-S. Chin, W. Chow, L. G. de Lima, P. C. Dishuck, R. Durbin, T. Dvorkina, I. T. Fiddes, G. Formenti, R. S. Fulton, A. Fungtammasan, E. Garrison, P. G. S. Grady, T. A. Graves-Lindsay, I. M. Hall, N. F. Hansen, G. A. Hartley, M. Haukness, K. Howe, M. W. Hunkapiller, C. Jain, M. Jain, E. D. Jarvis, P. Kerpedjiev, M. Kirsche, M. Kolmogorov, J. Korlach, M. Kremitzki, H. Li, V. V. Maduro, T. Marschall, A. M. McCartney, J. McDaniel, D. E. Miller, J. C. Mullikin, E. W. Myers, N. D. Olson, B. Paten, P. Peluso, P. A. Pevzner, D. Porubsky, T. Potapova, E. I. Rogaev, J. A. Rosenfeld, S. L. Salzberg, V. A. Schneider, F. J. Sedlazeck, K. Shafin, C. J. Shew, A. Shumate, Y. Sims, A. F. A. Smit, D. C. Soto, I. Sović, J. M. Storer, A. Streets, B. A. Sullivan, F. Thibaud-Nissen, J. Torrance, J. Wagner, B. P. Walenz, A. Wenger, J. M. D. Wood, C. Xiao, S. M. Yan, A. C. Young, S. Zarate, U. Surti, R. C. McCoy, M. Y. Dennis, I. A. Alexandrov, J. L. Gerton, R. J. O’Neill, W. Timp, J. M. Zook, M. C. Schatz, E. E. Eichler, K. H. Miga, A. M. Phillippy, The complete sequence of a human genome. Science 376, 44–53 (2022).

51. D. Li, R. Luo, C.-M. Liu, C.-M. Leung, H.-F. Ting, K. Sadakane, H. Yamashita, T.-W. Lam, MEGAHIT v1.0: A fast and scalable metagenome assembler driven by advanced methodologies and community practices. Methods 102, 3–11 (2016).

52. D. Li, C.-M. Liu, R. Luo, K. Sadakane, T.-W. Lam, MEGAHIT: an ultra-fast single-node solution for large and complex metagenomics assembly via succinct de Bruijn graph. Bioinformatics 31, 1674–1676 (2015).

53. P. Danecek, J. K. Bonfield, J. Liddle, J. Marshall, V. Ohan, M. O. Pollard, A. Whitwham, T. Keane, S. A. McCarthy, R. M. Davies, H. Li, Twelve years of SAMtools and BCFtools. Gigascience 10, giab008 (2021).

54. D. D. Kang, F. Li, E. Kirton, A. Thomas, R. Egan, H. An, Z. Wang, MetaBAT 2: an adaptive binning algorithm for robust and efficient genome reconstruction from metagenome assemblies. PeerJ 7, e7359 (2019).

55. D. H. Parks, M. Imelfort, C. T. Skennerton, P. Hugenholtz, G. W. Tyson, CheckM: assessing the quality of microbial genomes recovered from isolates, single cells, and metagenomes. Genome Res 25, 1043–1055 (2015).

56. M. R. Olm, C. T. Brown, B. Brooks, J. F. Banfield, dRep: a tool for fast and accurate genomic comparisons that enables improved genome recovery from metagenomes through de-replication. ISME J 11, 2864–2868 (2017).

57. P.-A. Chaumeil, A. J. Mussig, P. Hugenholtz, D. H. Parks, GTDB-Tk: a toolkit to classify genomes with the Genome Taxonomy Database. Bioinformatics 36, 1925–1927 (2020).

58. C. Hong, S. Manimaran, Y. Shen, J. F. Perez-Rogers, A. L. Byrd, E. Castro-Nallar, K. A. Crandall, W. E. Johnson, PathoScope 2.0: a complete computational framework for strain identification in environmental or clinical sequencing samples. Microbiome 2, 33 (2014).

59. E. A. Franzosa, L. J. McIver, G. Rahnavard, L. R. Thompson, M. Schirmer, G. Weingart, K. S. Lipson, R. Knight, J. G. Caporaso, N. Segata, C. Huttenhower, Species-level functional profiling of metagenomes and metatranscriptomes. Nat Methods 15, 962–968 (2018).

60. R Core Team, R: A Language and Environment for Statistical Computing (2022) (available at https://www.R-project.org/).

61. E. Paradis, K. Schliep, ape 5.0: an environment for modern phylogenetics and evolutionary analyses in R. Bioinformatics 35, 526–528 (2019).

62. J. Oksanen, F. G. Blanchet, M. Friendly, R. Kindt, P. Legendre, D. McGlinn, P. R. Minchin, R. B. O’Hara, G. L. Simpson, P. Solymos, M. H. H. Stevens, E. Szoecs, H. Wagner, vegan: Community Ecology Package (2020; https://CRAN.R-project.org/package=vegan).

63. D. Bates, M. Mächler, B. Bolker, S. Walker, Fitting Linear Mixed-Effects Models Using lme4. Journal of Statistical Software 67, 1–48 (2015).

64. A. Kuznetsova, P. B. Brockhoff, R. H. B. Christensen, lmerTest Package: Tests in Linear Mixed Effects Models. Journal of Statistical Software 82, 1–26 (2017).

65. R. D. Peng, simpleboot: Simple Bootstrap Routines (2019) (available at https://cran.r-project.org/web/packages/simpleboot/index.html).

66. A. C. Davison, D. V. Hinkley, *Bootstrap Methods and their Application* (Cambridge University Press, Cambridge, 1997; https://www.cambridge.org/core/books/bootstrap-methods-and-their-application/ED2FD043579F27952363566DC09CBD6A).

67. A. Canty, B. R. (author of parallel support), boot: Bootstrap Functions (Originally by Angelo Canty for S) (2022) (available at https://cran.r-project.org/web/packages/boot/index.html).

