## Supplemental Files for "Genotype-environment-driven dysbiosis in the skin microbiome of ichthyosis"

**Fig. S1. Severity and quality of scale vary in ichthyosis.** (**A**) Skin with an *ABCA12* mutation usually has small flaky scales with background erythema. This phenotype was formally known as congenital ichthyosiform erythroderma. Skin with a *TGM1* mutation usually has large, plate-like scales and mild erythema. This phenotype was formerly known as lamellar ichthyosis. Skin with a *KRT10* mutation usually has cobble-stone-like scale in a corrugated pattern and mild erythema. This phenotype is characteristic of epidermolytic ichthyosis. (**B**) Shins of five patients with *TGM1* mutation demonstrating the varying severity scale.

**Fig. S2. phylogenetic relationships of the 907 MAGs used for the analysis.**

**Supplementary Tables:**

https://www.dropbox.com/scl/fi/fys2urkmai0kghis8lob7/SupplementaryTables.xlsx?rlkey=7xsu1v1dh5vygpctgomiriksh&dl=0

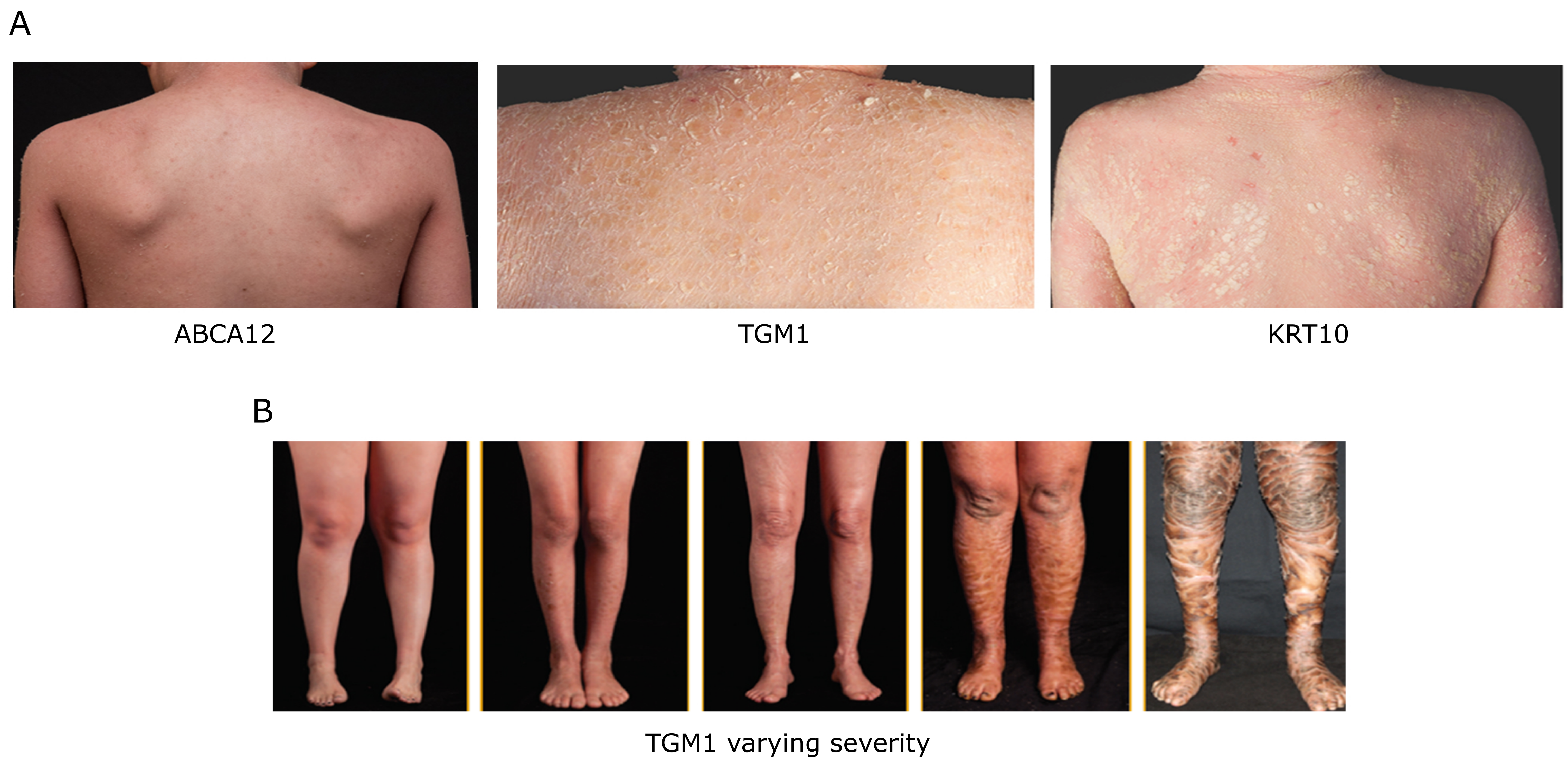


Fig S1

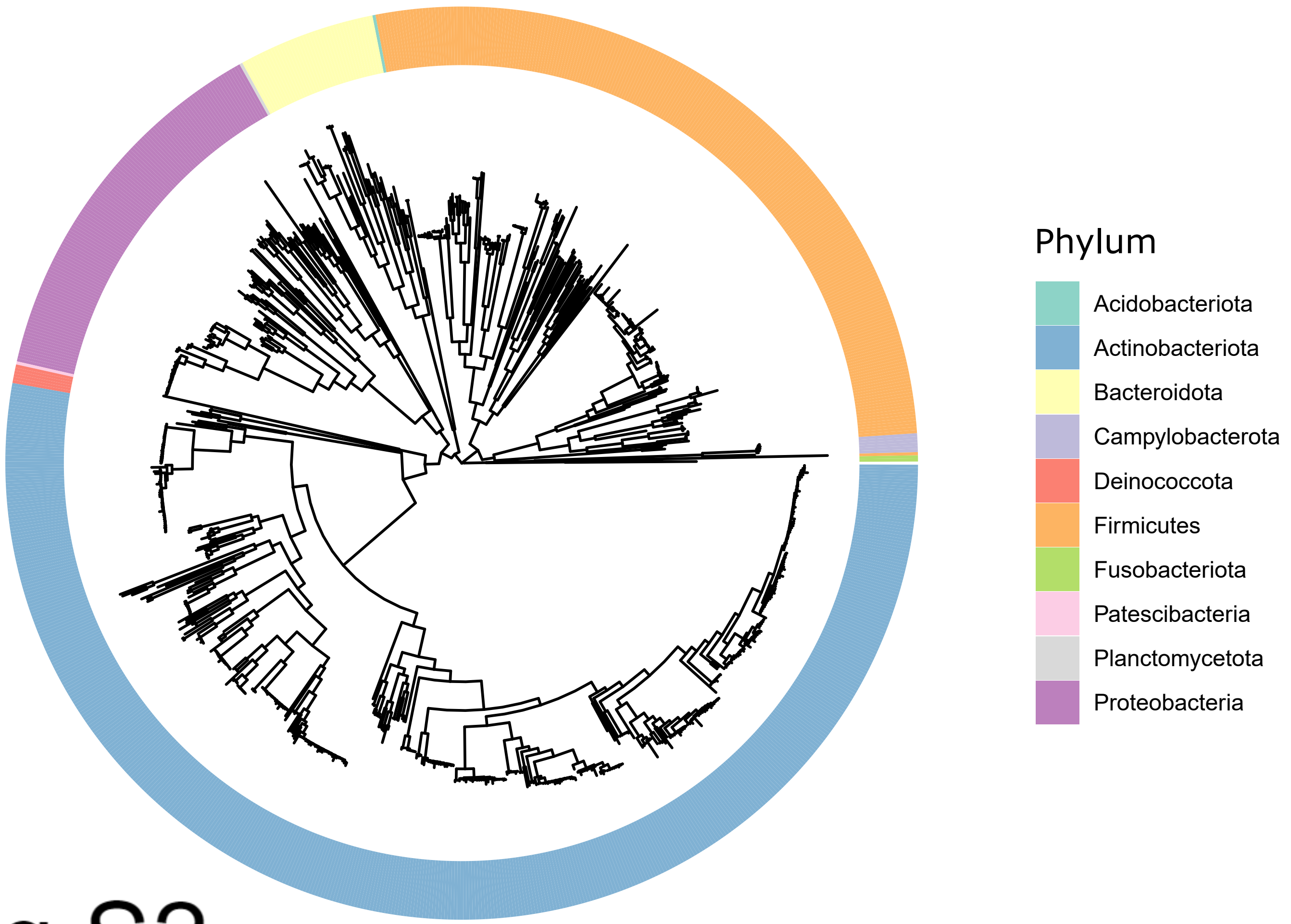
Phylum
Acidobacteriota
Actinobacteriota
Bacteroidota
Campylobacterota
Deinococcota
Firmicutes
Fusobacteriota
Patescibacteria
Planctomycetota
Proteobacteria

Fig S2

### Supplementary Note

Here we investigated skin microbiome signatures at the species, strain, and metabolic pathway levels in five ichthyosis genotypes (KRT2, KRT10, ABCA12, ALOX12B, and NIPAL4). Due to the rarity of these genotypes, we based the bulk of our analyses on grouping the genotypes by their direct effect on critical components of the stratum corneum: keratinopathic ichthyoses (KPI) affect the production of keratin filaments and are represented by KRT2 and KRT10, and lipid function ichthyoses (LFI) affect lipid metabolism/transfer functions and are represented by ABCA12, ALOX12B, and NIPAL4.

**Supplementary Note Tables:**

https://www.dropbox.com/scl/fi/rfd09pb747tdavwyito6g/SupplementaryNoteTables.xlsx?rlkey=io04t8ayw1zgzyzlglz9r999l&dl=0

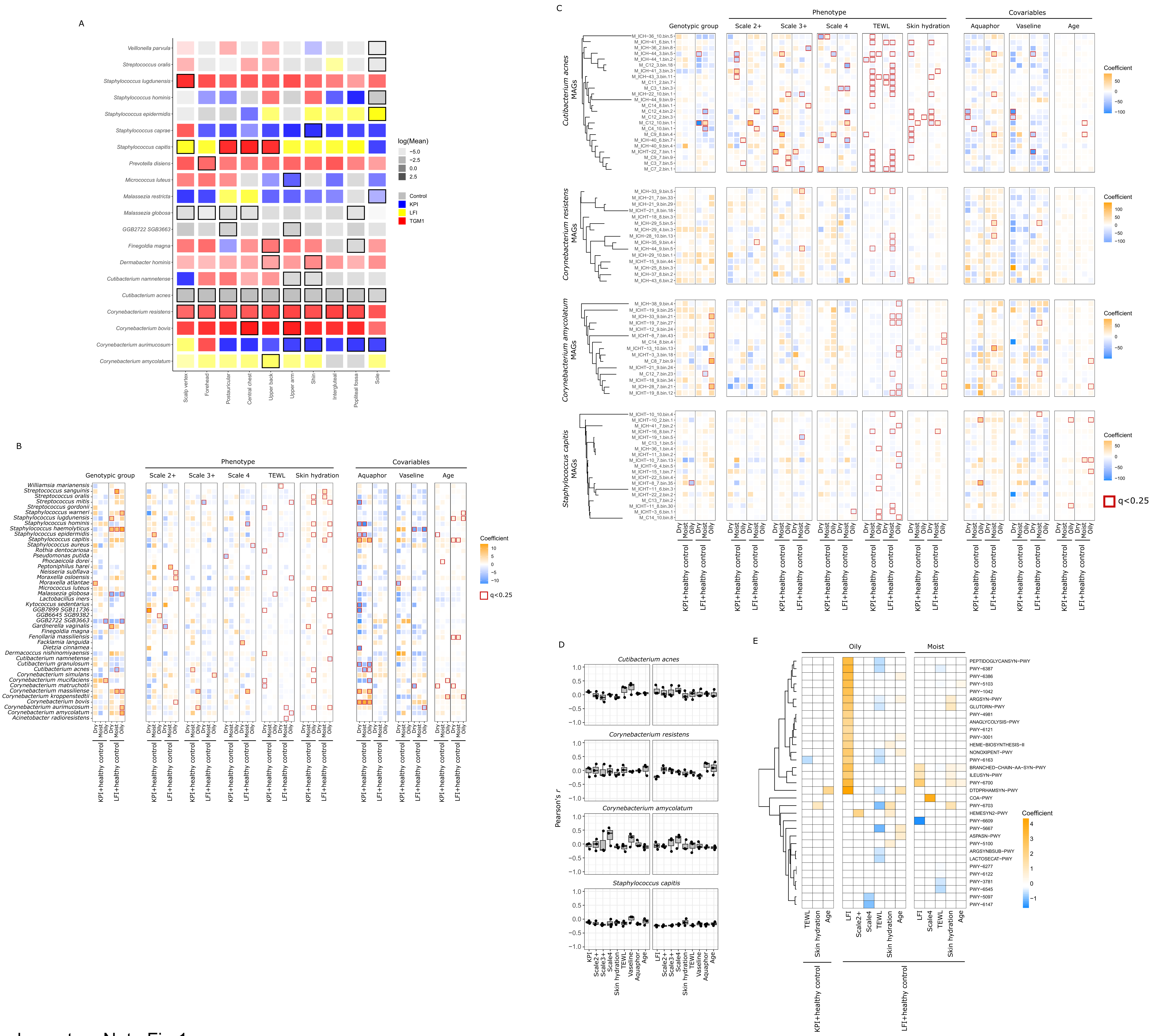


Supplementary Note Fig 1
